# Alzheimer’s disease-linked risk alleles elevate microglial cGAS-associated senescence and neurodegeneration in a tauopathy model

**DOI:** 10.1101/2024.01.24.577107

**Authors:** Gillian K. Carling, Li Fan, Nessa R. Foxe, Kendra Norman, Pearly Ye, Man Ying Wong, Daphne Zhu, Fangmin Yu, Jielin Xu, Allan Yarahmady, Hao Chen, Yige Huang, Sadaf Amin, Emmanouil Zacharioudakis, Xiaoying Chen, David M. Holtzman, Sue-Ann Mok, Evripidis Gavathiotis, Subhash C. Sinha, Feixiong Cheng, Wenjie Luo, Shiaoching Gong, Li Gan

## Abstract

The strongest risk factors for Alzheimer’s disease (AD) include the ε4 allele of apolipoprotein E (APOE), the *R47H* variant of triggering receptor expressed on myeloid cells 2 (TREM2), and female sex. Here, we combine *APOE4* and *TREM2R47H* (*R47H*) in female *P301S* tauopathy mice to identify the pathways activated when AD risk is the strongest, thereby highlighting disease-causing mechanisms. We find that the *R47H* variant induces neurodegeneration in female *APOE4* mice without impacting hippocampal tau load. The combination of *APOE4* and *R47H* amplified tauopathy-induced cell-autonomous microglial cGAS-STING signaling and type-I interferon response, and interferon signaling converged across glial cell types in the hippocampus. *APOE4-R47H* microglia displayed cGAS- and BAX-dependent upregulation of senescence, showing association between neurotoxic signatures and implicating mitochondrial permeabilization in pathogenesis. By uncovering pathways enhanced by the strongest AD risk factors, our study points to cGAS-STING signaling and associated microglial senescence as potential drivers of AD risk.

## INTRODUCTION

Alzheimer’s disease (AD) is the most common type of late-onset dementia. The major pathologies in AD include amyloid-beta (Aβ) plaques, neurofibrillary tau tangles, vascular dysfunction, and neuroinflammation. Recent studies show that most genetic risk variants associated with late-onset AD are present in microglia, indicating that microglia may play a causal role in the disease^1–3^. During the chronic inflammation that occurs in response to AD pathology, microglia suppress synaptic plasticity, increase synaptic pruning, phagocytose neurons, and secrete neurotoxic factors that contribute to neuronal death^4–8^. In addition to worsening neuronal degeneration, chronic microglial activation can also drive pathological tau aggregation in AD^8^. Microglial activation amplifies tau fibrillization in tauopathy mouse models, inflammation activates kinases that phosphorylate tau, and microglial depletion halts tau propagation^9–13^. These studies point to a contributing role of microglia-driven inflammation in pathological tau and the progression of AD.

The strongest genetic risk factors for late-onset AD include the ε4 allele of *APOE* and the *R47H* point mutation in the TREM2 receptor^14–17^. TREM2 is a single-pass transmembrane receptor expressed in myeloid lineage cells and found exclusively in microglia in the brain^18^. Microglial TREM2 can exert either protective or detrimental effects in AD depending on disease stage and context. TREM2 allows microglia to surround and compact Aβ plaques through APOE binding, subsequently reducing amyloid-mediated neuronal toxicity^3,19–21^. However, TREM2 activation could be detrimental in the context of tauopathy, likely through exacerbation of microglial activation^22,23^.

The rare *TREM2R47H* (*R47H*) variant increases AD risk 2- to 4.5-fold, making it second to *APOE4* in increasing AD probability^14,15^. *R47H* is associated with earlier AD onset^24^, faster cognitive decline^25,26^, and increased tau pathology^27,28^. *R47H* has been shown to worsen microglial inflammation^29,30^ and impair ligand binding^31–33^, weakening microglial ability to surround and compact Aβ plaques^20,34^, although further work is needed to characterize its deleterious effects. Notably, APOE is a ligand of microglial TREM2, and the *R47H* risk variant impairs APOE binding, potentially contributing to its disease risk^20,31,32,35,36^.

*APOE4* is the strongest and most common genetic risk factor for late-onset AD, where heterozygosity increases disease risk by 4-fold relative to two copies of *APOE* ε3 (*APOE3*), a similar odds ratio as *R47H*, and homozygosity for *APOE4* increases risk by up to 12-fold^16,17,37^. The *APOE* ε2 allele is protective against the development of AD^37^ relative to two copies of *APOE3*. *APOE4* is associated with earlier AD onset^17,26^, higher Aβ and tau toxicity^38,39^, and increased microglial inflammation^40–43^. Microglial depletion in humanized *APOE4* tauopathy mice blocks the toxic effects of the ε4 allele and halts neurodegeneration, suggesting that *APOE4* may exert its neurotoxic effects specifically through microglia^44^.

Although *APOE* is highly expressed in astrocytes, *APOE* is also strongly induced in microglia downstream of TREM2 activation in the context of disease^45,46^. Namely, TREM2 is required for the induction of a disease-associated microglia (DAM) signature^45^ and microglial neurodegenerative phenotype (MGnD)^47^ in response to disease pathology, signatures which both include *APOE* upregulation. TREM2 activation by Aβ pathology or apoptotic cells leads to downregulation of homeostatic gene expression and upregulation of *APOE* along with other pro-inflammatory genes^45,47^. It is currently unclear if this microglial state is protective or detrimental, since TREM2 deletion, and therefore blockage of this genetic signature, may be beneficial in reducing tau-mediated neurodegeneration^23,48^ but may worsen Aβ-dependent damage^34,45,49^. Conspicuously, the *R47H* risk variant increases microglial DAM induction and exacerbates *APOE* upregulation in the *P301S* tauopathy model, pointing to interplay between these two strong genetic risk factors in microglia^29^. Few studies have examined the combination of *R47H* and *APOE4* risk alleles in humans, although one study has found that AD patient carriers of both *R47H* and *APOE4* had faster disease progression compared to *APOE4* carriers with common variant (CV) *TREM2*^26^. There is currently limited information regarding how the *TREM2-APOE* pathway ultimately contributes to AD risk, and downstream mechanisms of this pathway are unknown. Since microglial activation can be either beneficial or deleterious depending on the context, the combination of AD risk factors *R47H* and *APOE4* may help highlight the mechanisms worsening disease and contributing to increased risk in humans.

There are noteworthy sex differences in AD incidence and pathology^50,51^, such that a greater proportion of AD patients are women^52^, and *APOE4*-associated risk is stronger in women^53^. We accordingly focused our studies on female mice to examine the effects of combining three substantial risk factors: *R47H*, *APOE4*, and female sex. We chose to use the *P301S* tauopathy model for this study because tau pathology more closely aligns with cognitive decline than Aβ pathology in human patients^54^, previous studies have implicated microglial *APOE4* in tau-mediated neurodegeneration^39,44^, and *R47H* has been found to upregulate microglial *APOE* in the *P301S* model^29^. We find that the combination of *APOE4* and *R47H* (*APOE4-R47H*) risk factors induces neurodegeneration and exacerbates microgliosis in female tauopathy mice. *APOE4-R47H* amplifies microglial cGAS-STING signaling and downstream type I interferon (IFN-I) response, as well as increasing microglial senescence. Mitochondrial BAX macropore inhibition or cGAS inhibition was sufficient to ameliorate microglial senescence, implicating mitochondrial membrane permeabilization and cGAS activation as involved in *APOE4-R47H* induction of senescence. Our study identifies microglial cGAS-STING activation and associated senescence as central mechanisms of AD, highlighting these pathways as critical therapeutic targets.

## RESULTS

### *APOE4-R47H* induces neurodegeneration and synaptic loss in female tauopathy mice

*APOE4* elevates AD risk in both males and females, with females exhibiting stronger effects^53^. To assess the effects of *APOE4-R47H* (*E4/E4/R47H/+*) on tauopathy-induced neurodegeneration, we first quantified body weight change in 9–10-month-old *P301S* transgenic versus non-transgenic female mice. Female mice experienced tauopathy-induced weight loss on both the *APOE3* and *APOE4* background (Figure 1A and S1A). *APOE4-R47H* exacerbated this tauopathy-induced weight loss compared to *APOE4-WT* (*E4/E4/mTrem2+/+*) (Figure 1A). We next measured plasma neurofilament light (NFL) concentration as a proxy measure for neurodegeneration^55,56^. *APOE4-R47H* exhibited significant increases in plasma NFL, whereas all other female tauopathy mice only showed trends towards elevated plasma NFL levels (Figure 1B and S1B).

**Figure 1.**
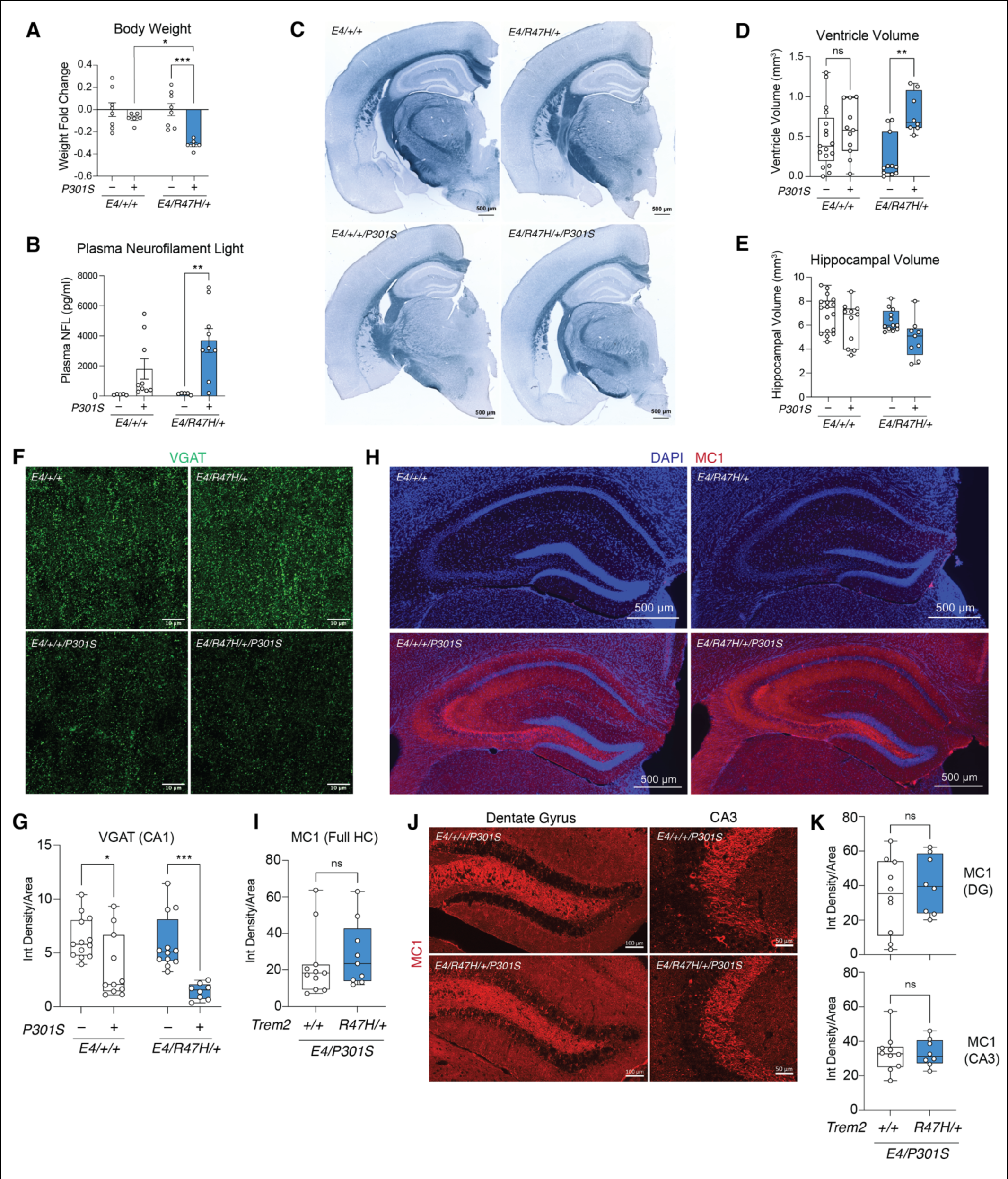
*APOE4-R47H* induces neurodegeneration and synaptic loss in female tauopathy mice (A) Quantification of body weight fold change in *P301S* transgenic versus non-transgenic female mice within each genotype at 9-10 months old. Each circle represents one animal. * p = 0.0223, *** p = 0.0009. Two-way ANOVA, Tukey’s multiple comparisons test. Data are reported as mean ± S.E.M. (B) Plasma neurofilament light concentration in female mice. Each circle represents one animal. ** p = 0.0086. Two-way ANOVA, Tukey’s multiple comparisons test. Data are reported as mean ± S.E.M. (C) Representative images of female mouse brain sections stained with Sudan black; scale bar, 500μm. (D and E) Quantification of ventricle volume (D) and hippocampal volume (E) based on staining from (C). Each circle represents sum volume measurements of five to seven brain sections per animal. Two statistically significant outliers were removed for ventricle volume. ** p = 0.0039, n.s. not significant. Two-way ANOVA, Tukey’s multiple comparisons test. Data are reported as boxplot with min. to max. (F) Representative immunofluorescence images of hippocampal CA1 striatum radiatum labeled with anti-VGAT (green); scale bar, 10μm. (G) Mean intensity of VGAT-positive puncta in the hippocampal CA1 striatum radiatum. Each circle represents mean intensity of three to four brain sections per animal. One statistically significant outlier was removed. * p = 0.0341, *** p = 0.0007. Two-way ANOVA, Tukey’s multiple comparisons test. Data are reported as boxplot with min. to max. (H) Representative immunofluorescence images of the full hippocampus labeled with anti-MC1 (red) and DAPI (blue); scale bar, 500μm. (I) Mean intensity of MC1 across the full hippocampus. Each circle represents mean intensity of three to four brain sections per animal. Two statistically significant outliers were removed. n.s. not significant. Unpaired two-tailed t-test. Data are reported as boxplot with min. to max. (J) Representative immunofluorescence images of hippocampal subregions labeled with anti-MC1 (red); scale bars, 100μm or 50μm, as labeled. (K) Mean intensity of MC1 within hippocampal subregions. Four statistically significant outliers were removed. n.s. not significant. Unpaired two-tailed t-test. Data are reported as boxplot with min. to max.

To directly examine the effects of *APOE4-R47H* on the brain, we quantified hippocampal and ventricular atrophy. Female tauopathy mice displayed mild neurodegeneration at 9-10 months old, with trends towards increased ventricle volume and decreased hippocampal volume (Figure 1C-E and S1C-E). *APOE4-R47H* significantly increased ventricle volume in female tauopathy mice, reflecting more severe brain atrophy than the other genotypes at this stage in disease (Figure 1C-D and S1C-D). To investigate these neurodegenerative effects of *APOE4-R47H* on a cellular level, we immunostained the hippocampal CA1 striatum radiatum of *APOE4* mice with an antibody for vesicular GABA transporter (VGAT) to quantify inhibitory neuronal synapses (Figure 1F). There was significant tauopathy-induced VGAT+ synapse loss in both *APOE4-WT* and *APOE4-R47H* mice, but synapse loss trended worse in *APOE4-R47H* mice (Figure 1F-G). Thus, *APOE4-R47H* tauopathy mice exhibited significant weight loss, increases in plasma NFL, brain atrophy, and synaptic loss (Figure 1A-G). In contrast, *R47H* had no discernable effect on body weight, plasma NFL, or brain atrophy on *APOE3* background in female tauopathy mice (Figure S1A-E).

Our previous studies showed that *R47H* exacerbated tau toxicity in female, but not male, *P301S* mice on the mouse *Apoe* background^29^. We next examined *APOE4-R47H* effects in males. Like female mice, male mice experienced tauopathy-induced weight loss and increases in plasma NFL at 9-10 months old (Figure S1F-G). Disease-associated weight loss trended worse in *APOE4-R47H* compared to *APOE4- WT* males (Figure S1F), but *APOE4-R47H* did not significantly affect plasma NFL concentration in male mice (Figure S1G). Consistent with previous studies^39^, male *APOE4-WT* mice displayed severe tauopathy-induced hippocampal atrophy (Figure S1H-J), more so than the atrophy observed in age-matched female tauopathy mice (Figure 1C-E). Intriguingly, *R47H* ameliorated hippocampal atrophy without affecting ventricle volume in male mice (Figure S1H-J). Thus, consistent with our sex-dimorphic effects of *R47H* on the mouse *Apoe* background, *R47H* exerts sex-dimorphic effects on the human *APOE4* background, accelerating neurodegeneration in females but ameliorating it in males. We focus our subsequent studies on mechanisms underlying *APOE4-R47H* effects in female tauopathy mice.

### *APOE4-R47H* does not significantly alter hippocampal tau load

We examined hippocampal tau load in 9–10-month-old female mice to determine if altered tau deposition underlies *APOE4-R47H* induction of neurodegeneration. Immunostaining with pathological conformation-specific tau antibody MC1 revealed no significant change in tau load across the hippocampus in *APOE4- R47H* compared to *APOE4-WT* tauopathy mice (Figure 1H-I). Additionally, no significant alterations in tau load induced by *APOE4-R47H* were detected in either the dentate gyrus or CA3 subregions (Figure 1J-K), indicating that *APOE4-R47H* mediated neurodegeneration is downstream of tau aggregation.

### *APOE4-R47H* expands an IFN-I microglial subcluster in the hippocampus

Microglia exhibit strong inflammatory responses in tauopathy^12,13^, and the TREM2 receptor is found exclusively in microglia in the brain^18^. Immunostaining for the microglia marker ionized calcium binding adaptor molecule 1 (IBA1) revealed tauopathy-induced microgliosis in the hippocampus, and *APOE4- R47H* significantly elevated tauopathy-provoked microgliosis compared to *APOE4-WT* (Figure 2A-B).

**Figure 2.**
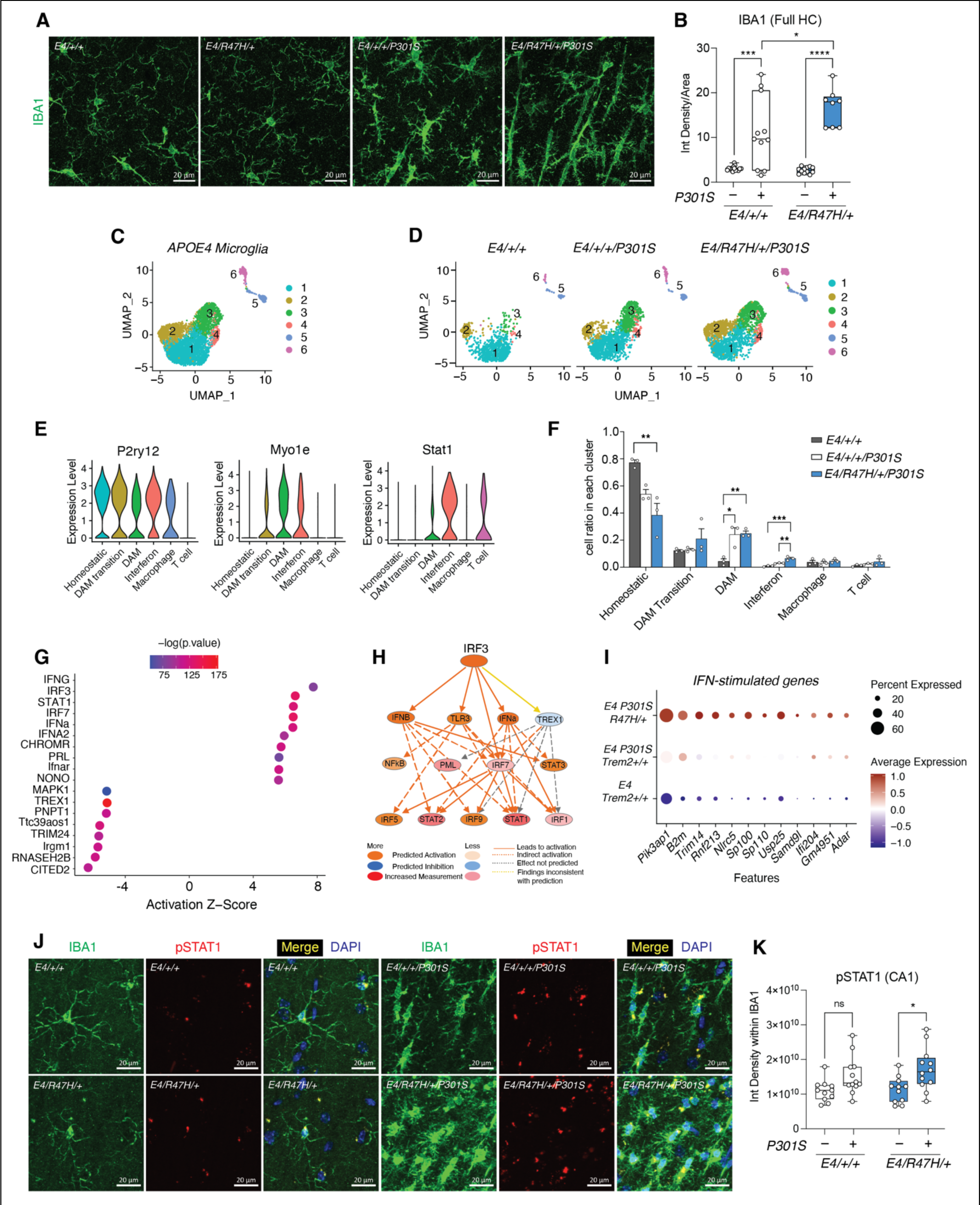
*APOE4*-*R47H* expands an IFN-I microglial subcluster in the hippocampus (A) Representative immunofluorescence images of the hippocampal CA1 subregion labeled with anti- IBA1 (green); scale bar, 20μm. B) Mean intensity of IBA1 across the full hippocampus. Each circle represents mean intensity of three to four brain sections per animal. One statistically significant outlier was removed. * p = 0.0374, *** p = 0.0002, **** p < 0.0001. Two-way ANOVA, Tukey’s multiple comparisons test. Data are reported as boxplot with min. to max. (C) UMAP plot of *APOE4* hippocampal immune cell nuclei colored according to subclusters. (D) UMAP plot of *APOE4* hippocampal immune cells split by genotype and colored according to subclusters. (E) Violin plots showing cell type markers split by immune cell subcluster to highlight cluster identity. (F) Quantification of cell ratio per microglial subcluster within each genotype. Microglial subcluster 4 is enriched in *APOE4-R47H*. Each circle represents one animal. * p < 0.05, ** p < 0.01, *** p < 0.001. Oneway ANOVA, Tukey’s multiple comparisons test. Data are reported as mean ± S.E.M. (G) Dot plot showing top IPA upstream regulator predictions for interferon signaling molecules based on microglial subcluster 4 markers, including IRF3 as a top activated upstream regulator. (H) IPA upstream regulator IRF3 activated network and its downstream predicted targets. (I) Dot plot showing interferon-stimulated genes enriched in microglial subcluster 4 and significantly upregulated in *E4/R47H/+/P301S* versus *E4/+/+/P301S* hippocampal microglia. (J) Representative immunofluorescence images of hippocampal CA1 subregion labeled with anti-IBA1 (green) and anti-pSTAT1 (red); scale bar, 20μm. (K) Quantification of pSTAT1 intensity within IBA1 overlapping regions in the hippocampal CA1 subregion. Each circle represents the mean quantification of three to four brain sections per animal. * p = 0.0269, n.s. not significant. Two-way ANOVA, Tukey’s multiple comparisons test. Data are reported as boxplot with min. to max.

To characterize the molecular signatures of microglial subpopulations affected by *APOE4-R47H*, we performed single-nuclei RNA sequencing (snRNA-Seq) on 9–10-month-old female mice. Quality control removed low-quality nuclei identified by total gene counts, unique molecular identifier (UMI) counts, and percentage of mitochondrial genes per nuclei. Sequencing reads from multiplets were excluded using DoubletFinder^57^. 91,744 nuclei passed quality control and were grouped into cell types through unsupervised clustering (Figure S2). The immune cell cluster, characterized by the expression of *Cx3cr1*, *P2ry12*, and *Csf1r* (Figure S2B), was further subclustered into six subpopulations in *APOE4* mice (Figure 2C-D). Subclusters 5 and 6 were identified as macrophage (*Mrc1*-high) and T cell (*Skap1*-high) clusters, respectively (Table S1). The remaining subclusters 1-4 were identified as microglia based on marker expression (Table S1). Microglial subcluster 1 contained high expression of homeostatic genes (*P2ry12*, *Cx3cr1*, *Selplg*, *Siglech*), whereas subcluster 3 was characterized by high levels of DAM signatures (*Myo1e*, *Cd83*, *Apoe*, *Spp1*) and reduced homeostatic gene expression (Figure 2E and Table S1). Subcluster 2 expressed intermediary levels of both homeostatic and DAM signatures, so this was deemed a cluster of microglia transitioning to the DAM state (Figure 2E and Table S1). In tauopathy mice, as expected, homeostatic microglial subcluster 1 was reduced and DAM subcluster 3 was significantly enriched (Figure 2F). *APOE4-R47H* did not affect the proportion of DAM microglia in subcluster 3 (Figure 2F).

Microglial subcluster 4 was highly enriched in IFN-I stimulated genes including *Stat1*, *Sp100*, *Trim30a*, and *Parp14* (Figure 2E and Table S1). Notably, this IFN-high subcluster 4 was significantly enriched in *APOE4-R47H* tauopathy microglia (Figure 2F). Ingenuity pathway analysis (IPA) of microglial subcluster 4 markers revealed IRF3 as a top upstream genetic regulator of this IFN signature (Figure 2G- H and Table S1). NF-κB and toll-like receptor signaling through TLR3 were among the other upstream regulators predicted downstream of IRF3 (Figure 2H and Table S1). *APOE4-R47H* microglia showed significant upregulation of tauopathy-induced IFN-stimulated genes present in the IFN-high subcluster 4 (Figure 2I and Table S2). To confirm *APOE4-R47H* exacerbation of IFN signatures on the protein level, we immunostained for phosphorylated STAT1 (pSTAT1) and found that indeed microglial pSTAT1 was significantly induced in the hippocampal CA1 subregion of *APOE4-R47H* tauopathy mice (Figure 2J-K).

We examined the effects of *R47H* on the *APOE3* background to determine if the exacerbation of microglial IFN signatures is driven primarily by *R47H*, or if it also requires *APOE4*. Immune cells subclustered into five groups on the *APOE3* background (Figure S3A-B). Based on marker expression, we characterized subcluster 1 as homeostatic microglia, subcluster 2 as DAM, subcluster 4 as macrophages, and subcluster 5 as T cells (Figure S3C and Table S3). Subcluster 3 had unclear identity with few cluster markers but included enrichment of complement genes (*C1qa*, *C1qc*) (Table S3). The only subcluster significantly impacted by *APOE3-R47H* (*E3/E3/R47H/+*) was subcluster 3, which was decreased in *P301S+* mice and was marginally restored by *APOE3-R47H* (Figure S3B). To determine if *APOE3-R47H* affects IFN response, we examined the expression levels of IFN-stimulated genes. IFN signature was induced in *APOE3* tauopathy mice, but *APOE3-R47H* did not significantly alter this effect (Figure S3D and Table S3). Our results show that the combination of *APOE4* and *R47H* risk factors significantly elevate microglial IFN-I response in the presence of tau pathology.

### *APOE4-R47H*-associated interferon signature converges across glial cell types

We next assessed the effects of *APOE4-R47H* on other glial cell types to see how they may be contributing to altered neurodegenerative phenotypes. Pseudobulk analyses were performed to compare *APOE4- R47H-P301S* versus *APOE4-WT-P301S* in oligodendrocyte and astrocyte populations. Pseudobulk analysis of oligodendrocytes showed immunoreactive markers (*C4b*, *Plin4*) among the top upregulated genes in *APOE4-R47H* (Figure 3A and Table S4). Notably, *APOE4-R47H* oligodendrocytes displayed upregulation of previously established disease-associated oligodendrocyte (DAO) markers^58^ including *C4b*, *H2-D1*, *H2-K1*, and *Plin4* (Figure 3B and Table S4). Pseudobulk analysis of astrocytes showed upregulation of metabolism-associated genes (*Phyhd1*, *Abca1*) and downregulation of NDMA receptor components (*Grin1, Grin2a*) in *APOE4-R47H* (Figure 3C and Table S5). *APOE4-R47H* astrocytes upregulated immunoreactive markers associated with disease response^59,60^, including several classic A1 markers *H2-D1*, *Ggta*, and *Fkbp5*, as well as pan-reactive markers *Gfap*, *C4b*, and *Stat3* (Figure 3D and Table S5).

**Figure 3.**
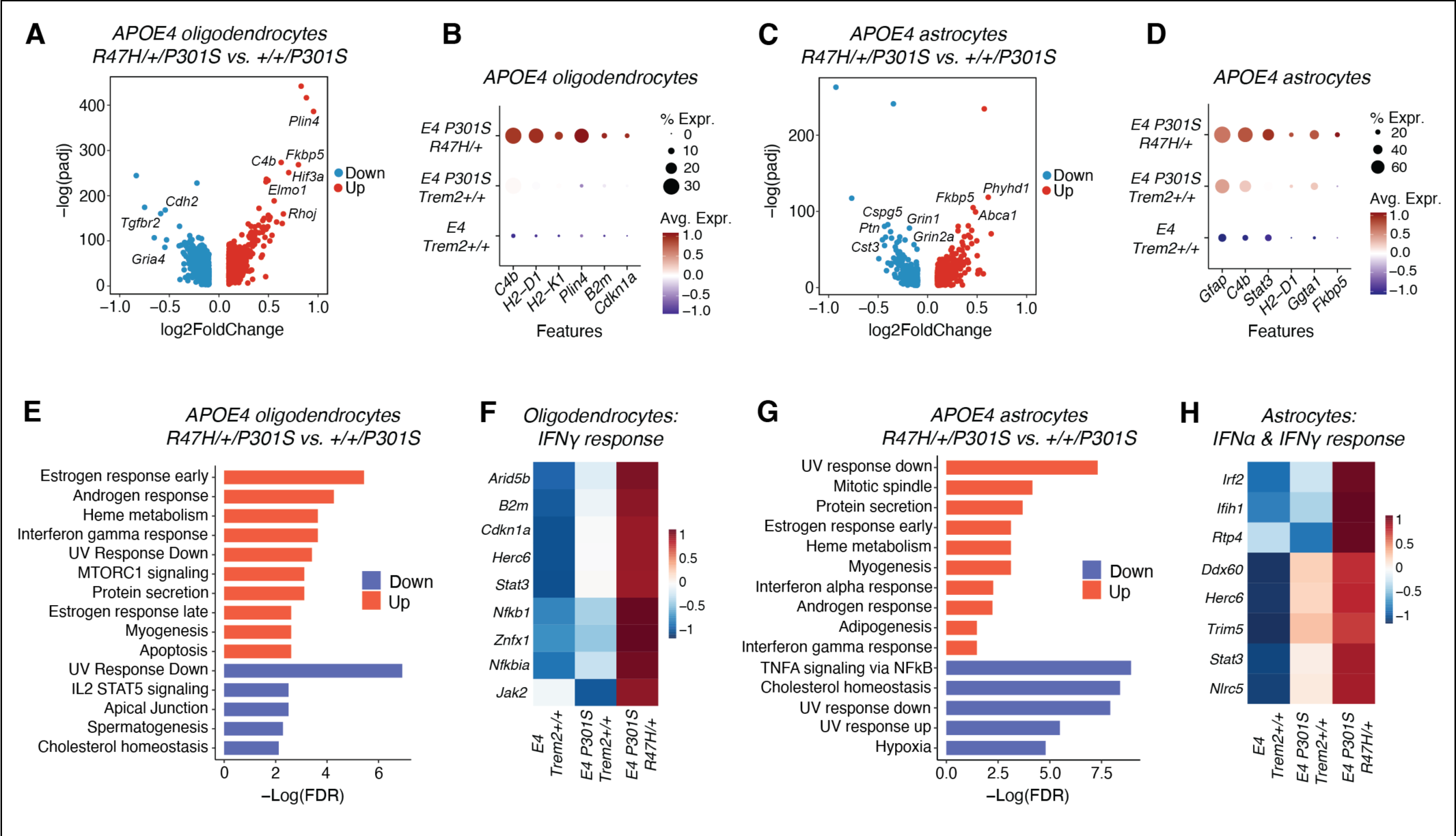
*APOE4-R47H*-associated IFN signature converges across glial cell types (A and C) Volcano plots of pseudobulk-seq data comparing *E4/R47H/+/P301S* versus *E4/+/+/P301S* hippocampal oligodendrocytes (A) and astrocytes (C) (n = 3 animals per genotype). Red and blue dots represent significant DEGs (p < 0.05). (B) Dot plot showing disease-associated oligodendrocyte genes significantly enriched in *E4/R47H/+/P301S* versus *E4/+/+/P301S* hippocampal oligodendrocytes. (D) Dot plot showing reactive astrocyte genes significantly enriched in *E4/R47H/+/P301S* versus *E4/+/+/P301S* hippocampal astrocytes. (E and G) Gene set enrichment analysis showing hallmark pathways based on the top 500 upregulated and downregulated pseudobulk-seq DEGs in *E4/R47H/+/P301S* versus *E4/+/+/P301S* hippocampal oligodendrocytes (E) and astrocytes (G). (F) Heatmap showing hallmark interferon gamma response genes that are significantly higher in *E4/R47H/+/P301S* versus *E4/+/+/P301S* hippocampal oligodendrocytes based on the pathway analysis in (E). (H) Heatmap showing hallmark interferon alpha and interferon gamma response genes that are significantly higher in *E4/R47H/+/P301S* versus *E4/+/+/P301S* hippocampal astrocytes based on the pathway analysis in (G).

Gene set enrichment analysis (GSEA) revealed upregulation of IFN-gamma and protein secretion hallmark pathways in *APOE4-R47H* tauopathy oligodendrocytes (Figure 3E and Table S4). Indeed, IFN-associated genes showed significant induction in *APOE4-R47H* oligodendrocytes compared to *APOE4- WT* oligodendrocytes in tauopathy mice (Figure 3F and Table S4). Consistent with the finding that IFN induces cell death in oligodendrocytes^61,62^, apoptosis was among the top hallmark pathways enriched in *APOE4-R47H* oligodendrocytes (Figure 3E and Table S4). In tauopathy astrocytes, GSEA highlighted upregulation of adipogenesis and downregulation of cholesterol homeostasis and hypoxia hallmark pathways in *APOE4-R47H* (Figure 3G and Table S5). Cholesterol metabolism pathways were also downregulated in *APOE4-R47H* oligodendrocytes (Figure 3E and Table S4), suggesting *APOE4-R47H* may contribute to metabolic dysfunction in glial cells. IFN-alpha, IFN-gamma, and protein secretion were additionally among the top upregulated pathways in *APOE4-R47H* astrocytes (Figure 3G and Table S5). IFN signatures were induced by tauopathy in *APOE4-WT* astrocytes, and this gene expression was significantly elevated by *APOE4-R47H* (Figure 3H and Table S5), similar to the effect seen in microglia and oligodendrocytes. Our results support a convergence of IFN signatures across glial cell types in *APOE4-R47H* tauopathy mice.

### *APOE4-R47H* exacerbates tauopathy-induced microglial cGAS-STING activation

Previous work in our lab has established that microglial cyclic GMP-AMP synthase (cGAS) drives IFN-I responses in the *P301S* tauopathy model^63^. cGAS is an antiviral DNA sensor that binds double-stranded DNA, catalyzing the synthesis of cyclic guanosine monophosphate-adenosine monophosphate (cGAMP) and activating stimulator of interferon genes (STING). This results in TBK1 phosphorylation, activating IRF3 and driving IFN-I production^64^. Given the strong IFN-I signatures in our *APOE4-R47H* tauopathy mice with IRF3 as a predicted upstream regulator (Figure 2F-I), we examined *Cgas* in our snRNA-Seq dataset and found that *Cgas* was a significant marker of the IFN-high microglial subcluster 4 (Figure 4A and Table S1). IPA predicted the cGAS-STING pathway as upstream of the IRF3-induced IFN-I response in *APOE4- R47H* microglia (Figure 4B and Table S1). To confirm *APOE4-R47H* upregulation of the cGAS-STING pathway on the protein level, we performed western blot for cGAS protein in hippocampal and frontal cortex mouse tissue lysates (Figure 4C-F). cGAS was induced by tauopathy in both *APOE4-R47H* and *APOE4- WT* hippocampi, but this response was only significant in the hippocampus of *APOE4-R47H* mice (Figure 4C and 4E). In the frontal cortex, tauopathy-induced cGAS upregulation was exclusively observed in *APOE4-R47H* and was significantly higher than in *APOE4-WT* tauopathy brains (Figure 4D and 4F).

**Figure 4.**
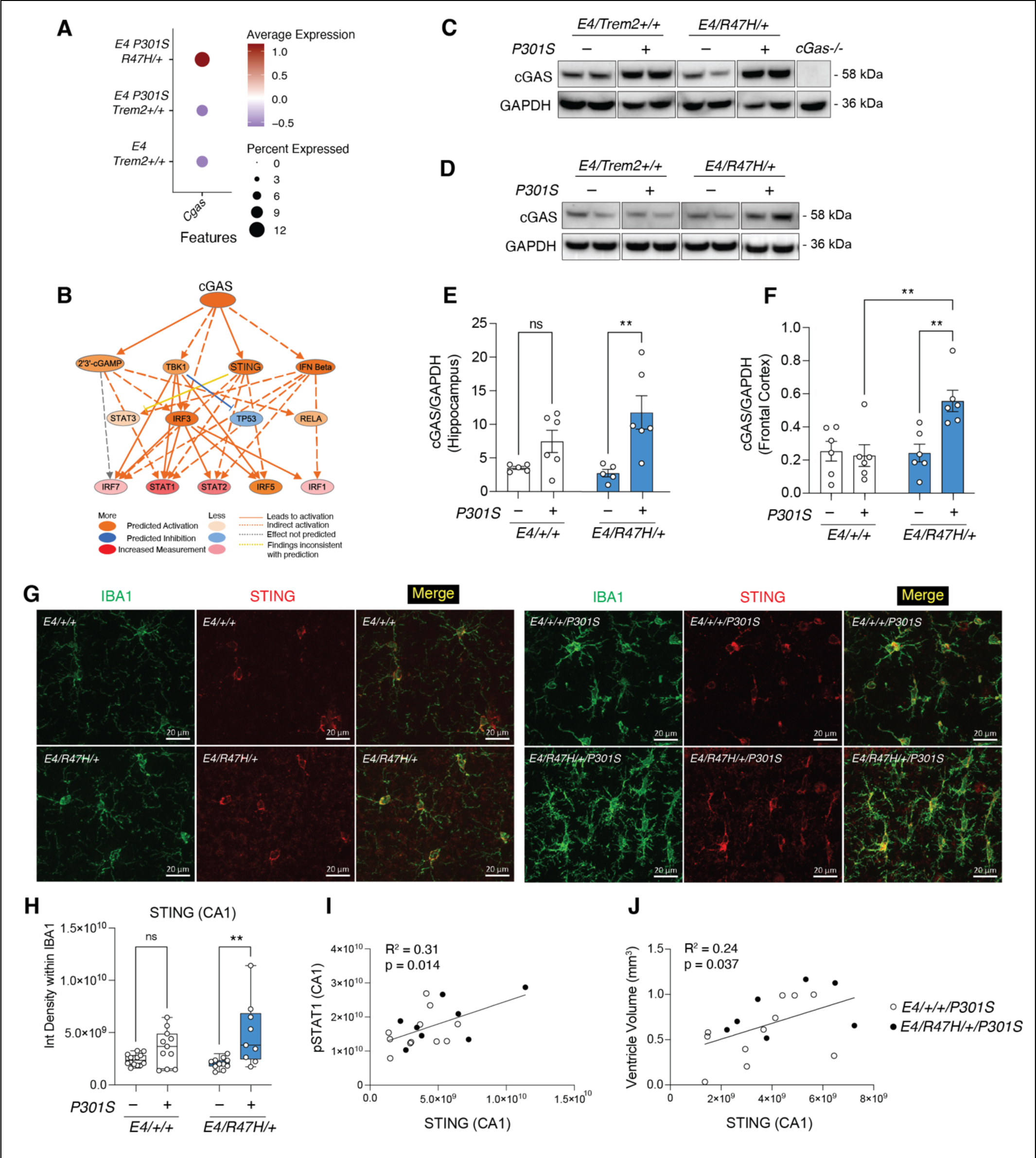
*APOE4*-*R47H* exacerbates tauopathy-induced microglial cGAS-STING activation (A) Dot plot showing *Cgas* expression, a microglial subcluster 4 marker, split by genotype. (B) IPA cGAS activated network and its downstream predicted targets. IPA predicts cGAS as an upstream regulator of microglial subcluster 4 (activation z-score = 3.573, p < 0.0001). (C and D) Representative western blots for cGAS and GAPDH using hippocampal (C) and frontal cortex (D) tissue lysates. (E and F) Western blot quantification of cGAS normalized to GAPDH in hippocampus (E) and frontal cortex (F). Each circle represents one animal. ** p < 0.01, n.s. not significant. Two-way ANOVA, Tukey’s multiple comparisons test. Data are reported as mean ± S.E.M. (G) Representative immunofluorescence images of hippocampal CA1 subregion labeled with anti-IBA1 (green) and anti-STING (red); scale bar, 20μm. (H) Quantification of STING intensity within IBA1 overlapping regions in the hippocampal CA1 subregion. Each circle represents the mean quantification of three to four brain sections per animal. ** p = 0.0022, n.s. not significant. Two-way ANOVA, Tukey’s multiple comparisons test. Data are reported as boxplot with min. to max. (I and J) Correlation scatterplot of microglial STING quantification compared to microglial pSTAT1 quantification (I) and brain ventricle volume quantification (J). Simple linear regression.

Since cGAS initiates an immune response via STING activation^64^, and cGAS-STING is primarily expressed by microglia in the brain^63,65,66^, we immunostained for STING in hippocampal CA1 microglia. STING was significantly induced by tauopathy in *APOE4-R47H* microglia but did not reach significance in *APOE4-WT* microglia (Figure 4G-H). Microglial STING levels significantly correlated with microglial pSTAT1 induction (Figure 4I), linking cGAS-STING upregulation with downstream IFN-I induction. Microglial STING also significantly correlated with ventricular atrophy (Figure 4J), suggesting this cGAS-STING induction either induces neurodegeneration or is stimulated in response to neuronal loss. Our previous research has found that cGAS-STING is a driver of synapse loss in the *P301S* model, and cGAS ablation rescues synapse loss and cognitive impairment^63^. Our results support that hyperactivation of cGAS-STING and IFN-I response could underlie the exacerbated neurodegeneration in *APOE4-R47H* tauopathy mice.

### *APOE4-R47H* microglial cGAS-STING induction is cell-autonomous

To further dissect *APOE4-R47H* effects on cGAS-STING and downstream pathways, we generated a new knock-in mouse line expressing the human *TREM2* common variant *CV/CV* or *R47H/CV* on the *APOE4/APOE4* homozygous background (hereby referred to as *APOE4-CV/CV* or *APOE4-R47H/CV*). Unlike our previous model, this new mouse line allows us to compare the *R47H* variant to human *TREM2- CV* instead of mouse *Trem2*. We confirmed correct recombination and insertion of *hTREM2-CV* and *hTREM2-R47H* at the *mTrem2* locus using polymerase chain reaction and Sanger sequencing (Figure S4). *TREM2* was expressed at similar levels in *R47H/CV* and *CV/CV* mouse brains, and mouse *Trem2* was not expressed (Figure S4B-C).

We isolated primary microglia from the new mouse line, treated with 0N4R tau fibrils, and performed bulk RNA-Seq to determine if the effects of the *R47H* mutation in primary microglia are comparable with those *in vivo* (Figure 5A). RNA-Seq exposed over 1,000 significant differentially expressed genes (DEGs) in *APOE4-R47H/CV* versus *APOE4-CV/CV* microglia after tau stimulation (Figure S5A and Table S6). In quality control, principal component analysis showed that samples cluster closely with their biological replicates, and sample-sample correlation revealed no outlier samples (Figure S5B-C). GSEA indicated *APOE4-R47H/CV* microglial enrichment of hallmark IFN pathways both at baseline (no stimulation) and after 0N4R tau fibril treatment (Figure 5B-C and Table S6). *Ifnb1* was one of the top DEGs in *APOE4- R47H/CV* versus *APOE4-CV/CV* microglia (Figure 5D and Table S6), and IPA predicted IFNB1 as an upstream regulator of other IFN signatures including IFNG, STAT1, and IRF1 (Figure 5E-F and Table S6). To establish if the cGAS-STING pathway was upregulated in *APOE4-R47H/CV* primary microglia, we ran western blot for cGAS in primary microglia lysates. Tau fibril stimulation strongly induced cGAS protein expression in *APOE4-R47H/CV* microglia, but not in *APOE4-CV/CV* microglia (Figure 5G-H). Therefore, cGAS and IFN enhancing effects of *APOE4-R47H* microglia were consistent in two models using either *mTrem2+/+* or *hTREM2-CV* as a comparison. Our findings further support cell-autonomous effects of cGAS and IFN upregulation induced by the *R47H* mutation on the *APOE4* background in tau-stimulated microglia.

**Figure 5.**
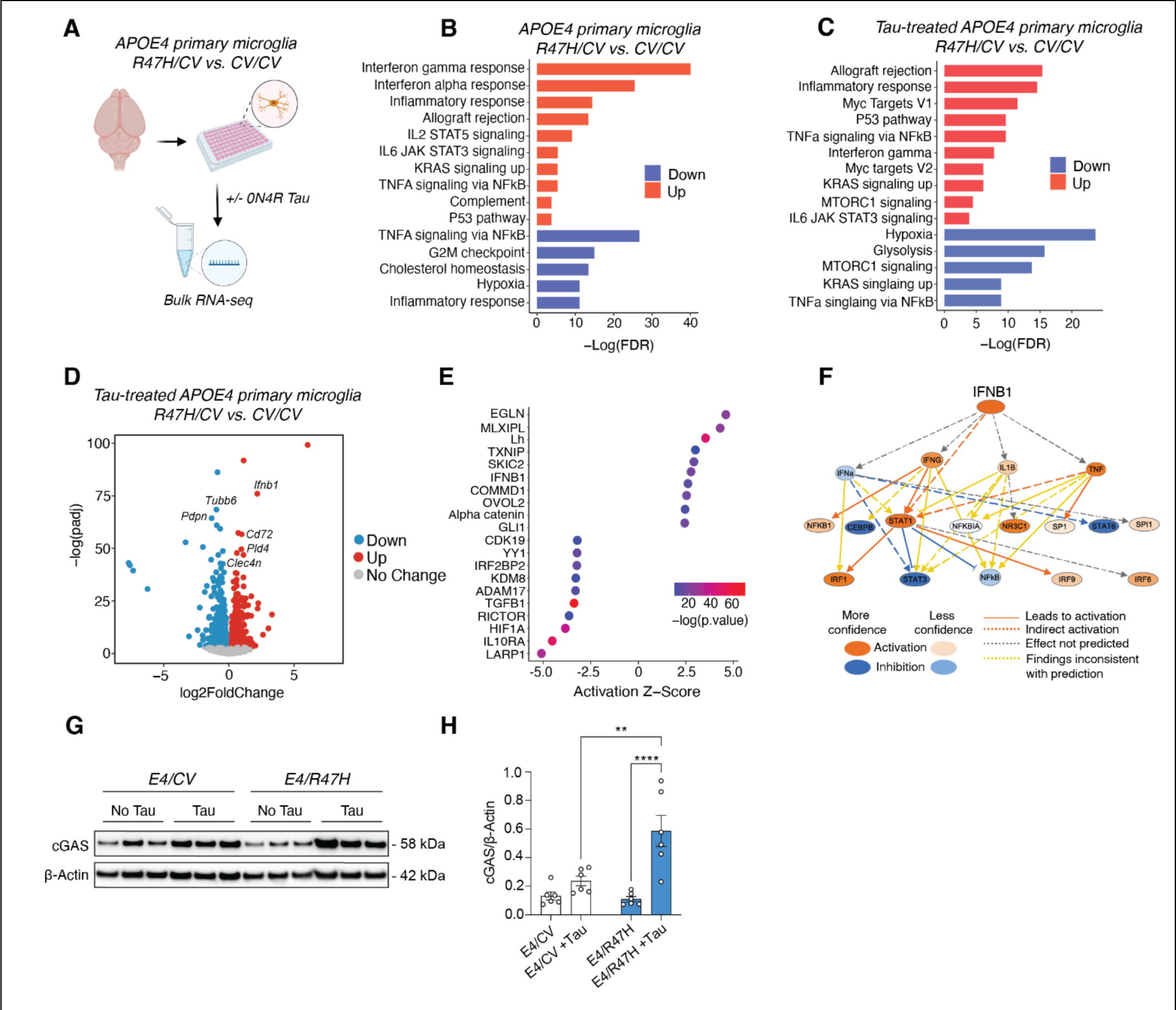
*APOE4*-*R47H* microglial cGAS-STING induction is cell-autonomous (A) Schematic showing bulk RNA-sequencing experimental design for *E4/CV* (*APOE4/APOE4; hTREM2-CV/CV*) and *E4/R47H (APOE4/APOE4; hTREM2-R47H/CV*) primary microglia at baseline and after 18-hour 0N4R tau fibril stimulation (n = 3 biological replicates per condition; n = 2 biologically independent experiments). Schematic created with BioRender.com. (B and C) Gene set enrichment analysis showing hallmark pathways based on the top 500 upregulated and downregulated DEGs in *E4/R47H* versus *E4/CV* primary microglia at baseline (B) and after tau fibril stimulation (C). (D) Volcano plot of RNA-seq data comparing *E4/R47H* versus *E4/CV* tau-stimulated primary microglia, revealing *Ifnb1* as a top DEG. Red and blue dots represent significant DEGs (p < 0.05). (E) Dot plot showing top IPA upstream regulator predictions for *E4/R47H* versus *E4/CV* tau-stimulated primary microglia, including IFNB1 as a top activated upstream regulator. (F) IPA upstream regulator IFNB1 activated network and its downstream predicted targets. (G) Representative western blots for cGAS and β-Actin using primary microglial lysates with or without 0N4R tau fibril stimulation. (H) Western blot quantification of cGAS/β-Actin. Each circle represents one biological replicate (n = 2 biologically independent experiments). ** p = 0.0025, **** p < 0.0001. Two-way ANOVA, Tukey’s multiple comparisons test. Data are reported as mean ± S.E.M.

### APOE4-R47H induces BAX- and cGAS-dependent senescence in tauopathy microglia

Tau pathology has recently been found to induce senescence in several cell types, including microglia, and clearance of these senescent cells ameliorates tau aggregation and cognitive decline^67–69^. Cellular senescence is characterized by permanent cell cycle arrest without cell death, along with induction of an inflammatory senescence-associated secretory phenotype (SASP)^70^. Notably, studies have linked cGAS-STING activation with senescence, finding that cGAS or STING inhibition suppresses SASP^71–75^. Since *APOE4-R47H* microglia have hyperactive cGAS-STING activation, we next determined if these cells also have altered senescence phenotypes.

To establish if senescence phenotypes are observed in our *in vivo* model, we first examined the expression of senescence-associated genes in *APOE4-R47H* versus *APOE4-WT* hippocampal mouse microglia. *APOE4-R47H* tauopathy microglia exhibited significant enrichment of senescence-associated genes compared to *APOE4-WT*, including *Ep300*, *B2m*, *Ikbkb*, and *Atm* (Figure 6A and Table S2). p21 is a cyclin-dependent kinase inhibitor that inhibits cell cycle progression and is frequently used as a marker of senescence^70^. Immunostaining revealed significant induction of senescence marker p21 in *APOE4- R47H* microglia, but not *APOE4-WT* microglia, in the presence of tau pathology (Figure 6B-C). Microglial p21 levels significantly correlated with hippocampal MC1-positive tau load, suggesting that higher tau burden could worsen senescence (Figure 6D). Microglial p21 levels also strongly correlated with microglial STING upregulation, supporting a positive association between aggravated cGAS-STING signaling and senescence (Figure 6E). In sum, *APOE4-R47H* induces tauopathy-associated microglial senescence in the hippocampus, and heightened p21 levels coincide with heightened cGAS-STING signaling *in vivo*.

**Figure 6.**
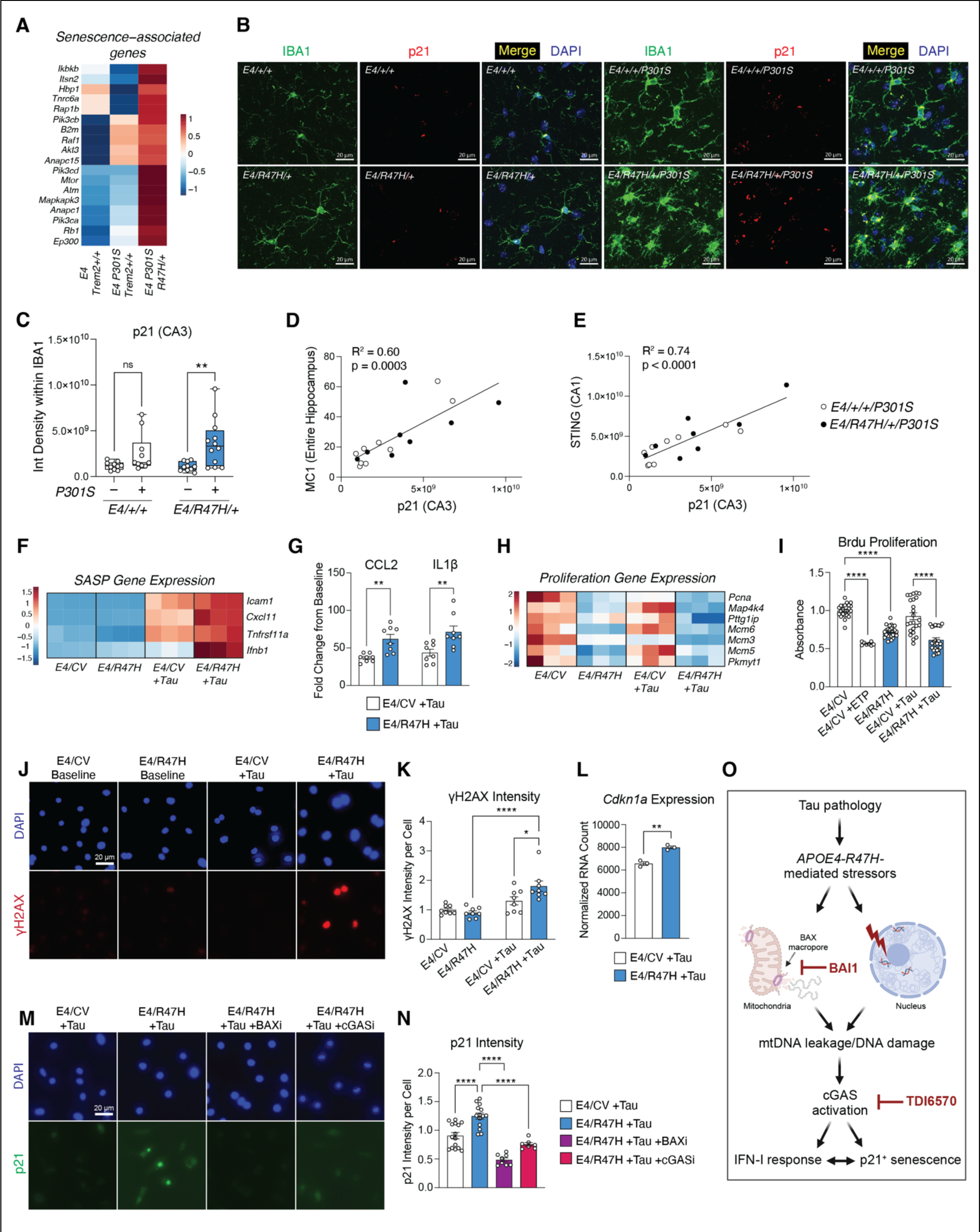
*APOE4*-*R47H* induces BAX- and cGAS-dependent senescence in tauopathy microglia (A) Heatmap showing senescence-associated genes that are significantly higher in *E4/R47H/+/P301S* versus *E4/+/+/P301S* hippocampal mouse microglia *in vivo*. (B) Representative immunofluorescence images of hippocampal CA3 subregion labeled with anti-IBA1 (green) and anti-p21 (red); scale bar, 20μm. (C) Quantification of p21 intensity within IBA1 overlapping regions in the hippocampal CA3 subregion. Each circle represents the mean quantification of three to four brain sections per animal. ** p = 0.0092, n.s. not significant. Two-way ANOVA, Tukey’s multiple comparisons test. Data are reported as boxplot with min. to max. (D and E) Correlation scatterplot of microglial p21 quantification compared to hippocampal MC1 quantification (D) and microglial STING quantification (E). Simple linear regression. (F) Heatmap showing senescence-associated secretory phenotype (SASP) genes that are significantly enriched in *E4/R47H* versus *E4/CV* tau-treated primary mouse microglia. (G) Bar plot showing fold change in microglial secretion of CCL2 and IL1β SASP cytokines after 18-hour 0N4R tau fibril stimulation. Data is normalized to cell count per well. Each circle represents one biological replicate (n = 2 biologically independent experiments). ** p < 0.01. Two-way ANOVA, Bonferroni multiple comparisons test. Data are reported as mean ± S.E.M. (H) Heatmap showing proliferation genes that are significantly diminished in *E4/R47H* versus *E4/CV* tau-treated primary mouse microglia. (I) Bar plot showing Brdu proliferation ELISA in primary microglia at baseline, after 18-hour etoposide exposure, or after 18-hour 0N4R tau fibril stimulation. Each circle represents one biological replicate (n = 3 biologically independent experiments, except etoposide n = 1 experiment). **** p < 0.0001. One-way ANOVA, Tukey’s multiple comparisons test. Data are reported as mean ± S.E.M. (J) Representative immunofluorescence images of primary microglia labeled with anti-γH2AX (red) and DAPI (blue) after 72-hour 0N4R tau fibril stimulation; scale bar, 20μm. (K) Mean γH2AX intensity per cell in primary microglia after 72-hour 0N4R tau fibril exposure. Each circle represents mean intensity from six images per biological replicate (n = 2 biologically independent experiments). * p = 0.0298, **** p < 0.0001. Two-way ANOVA, Tukey’s multiple comparisons test. Data are reported as mean ± S.E.M. (L) Bar plot showing *Cdkn1a* normalized RNA counts after 18-hour 0N4R tau fibril stimulation in primary microglia. Each circle represents one biological replicate. ** p = 0.0021. Unpaired two-tailed t-test. Data are reported as mean ± S.E.M. (M) Representative immunofluorescence images of primary microglia labeled with anti-p21 (green) and DAPI (blue) after 72-hour 0N4R tau fibril exposure, with or without BAX inhibition (BAI1) and cGAS inhibition (TDI6570); scale bar, 20μm. (N) Mean p21 intensity per cell in primary microglia after 72-hour 0N4R tau fibril exposure, with or without BAX inhibition (BAI1) and cGAS inhibition (TDI6570). Each circle represents mean intensity from six to eight images per biological replicate (n = 4 biologically independent experiments, except BAXi and cGASi n = 2 experiments). **** p < 0.0001. One-way ANOVA, Tukey’s multiple comparisons test. Data are reported as mean ± S.E.M. (O) Working model illustrating proposed mechanism of senescence induction in *APOE4-R47H* tauopathy microglia. Tau and *APOE4-R47H*-related mitochondrial stressors may increase BAX-mediated mitochondrial membrane permeabilization, facilitating the release of mtDNA and increasing DNA damage. mtDNA release and DNA damage activate cGAS, which upregulates IFN-I response and senescence. Inhibition of BAX or cGAS is sufficient to block p21^+^ senescence. Schematic created with BioRender.com.

We further dissected the *APOE4-R47H* microglial senescence phenotype in primary microglia. *APOE4-R47H/CV* primary microglia significantly enriched the p53 tumor suppressor pathway both at baseline and after tau fibril stimulation (Figure 5B-C and Table S6), which can activate p21, resulting in cell cycle arrest and induction of senescence^76,77^. Examination of our bulk RNA-Seq dataset further revealed exaberbation of SASP gene expression in tau fibril-stimulated *APOE4-R47H/CV* primary microglia, including *Icam1*, *Cxcl11*, and *Tnfsf11a* (Figure 6F and Table S6). Measurement of cytokine secretion in microglial conditioned media confirmed that *APOE4-R47H/CV* primary microglia secrete significantly more CCL2 and IL1β than *APOE4-CV/CV* upon stimulation with tau fibrils (Figure 6G). Both CCL2 and IL1β production can be stimulated by cGAS and IFN-I response^64,78,79^, implicating cGAS as a driver of the observed SASP signature.

Senescence is associated with cell cycle arrest. *APOE4-R47H/CV* primary microglia showed significant reduction in genes related to proliferation, including *Pcna*, *Map4k4*, and *Mcm5*, both at baseline (unstimulated) and after tau fibril stimulation (Figure 6H and Table S6). We validated this phenotype by measuring proliferation using 5-bromo-2’-deoxyuridine (Brdu) incorporation in the primary microglia. Microglial proliferation trended lower after tau fibril stimulation in both *APOE4-R47H/CV* and *APOE4- CV/CV*, suggesting tau fibrils may promote cell cycle arrest (Figure 6I). *APOE4-R47H/CV* reduced microglial proliferation to a similar degree as etoposide-treated *APOE4-CV/CV* microglia both at baseline (unstimulated) and after tau fibril stimulation (Figure 6I). Etoposide is a topoisomerase II inhibitor that generates double-stranded DNA breaks (DSBs), resulting in DNA damage and cellular senescence^80,81^, highlighting that the combination of *APOE4* and *R47H* risk alleles behaves similarly to a DNA damaging agent in arresting the microglial cell cycle. To directly assess this effect, we stained for γH2AX, which labels DSBs, as a readout for DNA damage, another hallmark of senescence^82,83^. Tau fibril stimulation led to significant γH2AX induction in *APOE4-R47H/CV* microglia, and tau-stimulated *APOE4-R47H/CV* microglia had significantly higher γH2AX compared to *APOE4-CV/CV* microglia (Figure 6J-K). Taken together, *APOE4-R47H/CV* tau-stimulated primary microglia display increased SASP expression and production, reduced proliferation, and increased DNA damage.

A recent study found that mitochondrial outer membrane permeabilization in a minority subset of mitochondria is a feature of cellular senescence^75^. This mitochondrial membrane permeabilization is mediated by BAX and BAK macropores, which facilitate mitochondrial DNA (mtDNA) release into the cytosol, triggering cGAS-STING activation and SASP^75^. We aimed to determine if BAX-dependent mitochondrial membrane permeabilization contributes to senescence in our model and if cGAS activation is required for this *APOE4-R47H/CV*-mediated senescence. RNA expression of *Cdkn1a*, the gene encoding senescence marker p21, was significantly increased in tau-stimulated *APOE4-R47H/CV* compared to *APOE4-CV/CV* primary microglia (Figure 6L and Table S6). We examined the protein levels of p21 after prolonged 72-hour tau fibril exposure in the presence of BAX inhibitor BAI1^84^ or cGAS inhibitor TDI6570^85^. Immunostaining revealed that *APOE4-R47H/CV* microglia amplify tau fibril-induced p21 signaling, and BAX inhibition was sufficient to abrogate p21 induction (Figure 6M-N). cGAS inhibition worked similarly as BAX inhibition to reduce p21, showing that both BAX activation and cGAS activation are required for *APOE4-R47H/CV* induction of p21+ senescence (Figure 6M-N).

Mitochondrial membrane permeabilization mediated by BAX can lead to both DNA damage^86^ and mtDNA leakage^75^. Numerous studies show that mtDNA leakage can induce cGAS activation and downstream SASP^63,66,71,75,87,88^, and we have previously found that tau fibrils induce mtDNA leakage in microglia^63^. In our model, BAX-mediated mitochondrial membrane permeabilization may lead to increased mtDNA leakage and DNA damage in *APOE4-R47H* microglia, contributing to aberrant cGAS-STING activation, which then in turn may drive the senescence phenotype through SASP and chronic interferon production^73,89–91^ (Figure 6O). In sum, our results support the model that *APOE4-R47H* microglia increase tauopathy-stimulated cGAS-STING signaling and IFN-I response, resulting in BAX- and cGAS-dependent senescence and accelerated neurodegeneration.

## DISCUSSION

*APOE4* and *R47H* are among the strongest genetic risk factors for late-onset AD, but mechanisms underlying disease risk and roles of the *TREM2-APOE* pathway in microglia are currently obscure. Our study has found that the combination of *APOE4* and *R47H* risk factors accelerates neurodegeneration in female tauopathy mice without significantly impacting hippocampal tau load. A previous study similarly found that *R47H* does not affect hippocampal tau load in the *P301S* model^29^, suggesting that *R47H* risk is driven by disease-enhancing microglial responses to tau, rather than by tau itself. Indeed, we find that *APOE4-R47H* exacerbates microgliosis, amplifies microglial cGAS-STING signaling and downstream IFN response pathways in a cell-autonomous manner, and increases cGAS- and BAX-dependent microglial senescence. Our study links the strongest AD risk factors, *R47H*, *APOE4*, and female sex, with microglial cGAS-STING activation and associated senescence in a tauopathy model, highlighting these pathways as important therapeutic targets in the treatment of AD and primary tauopathies.

Female mice exhibited milder tauopathy-induced neurodegeneration than male mice at 9-10- months-old, such that females only showed trends towards ventricular and hippocampal atrophy, whereas males displayed severe, significant atrophy of the hippocampus. This sex-dependent effect in the *P301S* model has been previously reported, such that male *APOE4-P301S* mice experience higher levels of hippocampal and entorhinal/piriform cortex atrophy than *APOE4-P301S* female mice^92^. In our model, the combination of *APOE4* and *R47H* risk alleles was sufficient to induce neurodegeneration and synaptic loss in 9-10-month-old female tauopathy mice, whereas *APOE4* alone and *APOE3-R47H* did not yet exhibit significant neurodegeneration. This suggests the combination of *APOE4* and *R47H* in tauopathy mice accelerates disease progression in females, inducing neurodegeneration at a younger age. Remarkably, the combination of *APOE4* and *R47H* risk alleles in male mice had the opposite effect, ameliorating the severe hippocampal neurodegeneration seen with *APOE4* alone. A previous study using only male *APOE4-P301S* mice found that the absence of *TREM2* worsened neurodegeneration and increased a subset of tau markers without affecting MC1+ tau^93^, similar to our findings reported here in female *APOE4- R47H* mice. The *R47H* risk variant, however, was not studied in that work^93^, and *R47H* actually had protective effects in our male *APOE4-P301S* mice, showing that *R47H* does not necessarily behave like *TREM2* knockout. Similar to our results, a previous study using male mice found *R47H* to be protective in the *P301S* model, reducing brain atrophy and microglial reactivity compared to *CV-TREM2*^94^. Consistent with these observations, our previous study found that *R47H* augments spatial memory deficits exclusively in female, but not male, *P301S* mice^29^. While the underlying mechanisms of this effect are currently unclear, previous studies have implicated the *TREM2-APOE* pathway. TREM2 induction of the MGnD signature, including upregulation of *APOE*, is sex-specific^47^, and *R47H* upregulation of the DAM signature, including *APOE* expression, is only seen in female tauopathy mice^29^. These studies emphasize the importance of characterizing *R47H* in sex-stratified studies, a critical topic to be addressed in future research.

cGAS-STING and downstream IFN signaling are likely drivers of microglia-mediated neurodegeneration in female *APOE4-R47H* mice, as this pathway has been previously implicated as a causal factor in neurodegeneration. Inhibition of cGAS, STING, or IFN is protective against microgliosis and neurodegeneration in several different models of aging and AD^63,71,95^, and STING activation is sufficient to induce neurodegeneration and cognitive decline in the absence of AD pathology^71^. The cGAS promoter contains IFN response elements, so initial cGAS-induced IFN response can then stimulate further cGAS upregulation in a positive feedback loop, contributing to neuroinflammation^96^. Frontotemporal dementia patients show upregulation of viral defense response genes, including IFN-I, and sporadic AD patients carrying *R47H* upregulate IFN-I along with other pro-inflammatory cytokines, showing the relevance of this pathway to the human disease state^30,97^. Intriguingly, a recent study found that activated microglia recruit T cell infiltration in the brains of *P301S* mice carrying *APOE4*, and this T cell infiltration was found to be a driver of neurodegeneration in tauopathy^92^. In our model, amplified IFN signaling in *APOE4-R47H* tauopathy microglia could increase T cell infiltration, resulting in exacerbated neurodegeneration. Whether combining *APOE4* and *R47H* risk factors enhances T cell infiltration in female tauopathy brains remains to be seen.

Our study found that IFN-I upregulation converged across glial cell types in *APOE4-R47H* mouse brains. As microglia are the primary expressers of both TREM2 and cGAS-STING in the brain^18,63,65,66^, this IFN response likely begins in microglia and then propagates to other glial cell types. Microglia are known to crosstalk with other glial cells, as microglial activation is required to induce neurotoxic, activated astrocytes^60^, and microglial secreted factors impact oligodendrocyte capacity for remyelination^62^. Fittingly, *Cre*-induced cGAS activation in microglia results in upregulation of IFN-related transcripts in astrocytes and oligodendrocytes in the mouse brain^71^, mirroring the response seen in our *APOE4-R47H* brains. IFN exposure has been found to detrimentally impact oligodendrocytes, inhibiting oligodendrocyte progenitor cell (OPC) differentiation and inducing cell death in both oligodendrocytes and OPCs^61,98^. IFN-mediated oligodendrocyte damage may be one of the mechanisms through which cGAS-STING activation promotes neurodegeneration.

Recent studies have implicated cellular senescence in tauopathy pathogenesis^67–69^. Studies have identified cGAS as an integral component of cellular senescence, and it is well-established that cGAS or STING inhibition prevents SASP production in senescent cells^71–75^. Several studies have found that STING inhibition exclusively prevents SASP, but not other signatures of senescence, including p21, SA-β-Gal, and p16^71,74,75^. Interestingly, studies have found that cGAS deletion blocks these senescence markers (p21, SA-β-Gal, and p16) along with SASP, whereas STING deletion was less effective, indicating that cGAS and STING may play differing roles in senescence^72,73^. In our model, cGAS inhibition was sufficient to block p21 induction in *APOE4-R47H* primary microglia in the presence of pathological tau, further establishing the essential role of cGAS in the induction of senescence. It is possible that cGAS activation induces senescence through its upregulation of IFN-I response. Chronic IFNβ exposure stimulated by IRF3 triggers DNA damage, p53 pathway activation, and induction of senescence^73,89–91^. In turn, inhibition of the IFN-I pathway prevents senescence^73,89–91^. Further, IRF3 has been found to directly interact with BAX protein, affecting cell survival pathways through mitochondrial outer membrane permeabilization^99,100^, which is known to cause DNA damage^86^. IRF3 was predicted to be a top upstream regulator of the hippocampal microglial IFN subcluster enriched in *APOE4-R47H*, and the p53 pathway was enriched in *APOE4- R47H/CV* compared to *APOE4-CV/CV* primary microglia, implicating these pathways as potential cGAS-induced upstream mechanisms of *APOE4-R47H* microglial senescence. The effectiveness of BAX inhibition or cGAS inhibition at reducing senescence in *APOE4-R47H* microglia supports the causal role of cGAS stimulation and mitochondrial outer membrane permeabilization in the induction of senescence.

Mitochondrial stress-induced mtDNA release into the cytosol mediated by BAX and BAK macropores has been identified as a mechanism by which cGAS is activated in the context of aging, injury, and disease^63,66,71,75,87,88^. In a cell culture model, tau fibrils localized to the mitochondria and lysosomes of primary microglia, and tau treatment resulted in increased cytosolic mtDNA release, along with cGAS-STING activation^63^. AD risk factors *APOE4* and *R47H* are both associated with metabolic dysfunction, potentially rendering the mitochondria more vulnerable to tau-induced insult when combined. Accordingly, we observed changes to cholesterol homeostasis, adipogenesis, and hypoxia pathways in our *APOE4- R47H* glial cells. *APOE4* is known to disrupt glial lipid metabolism and alter the balance between mitochondrial and glycolytic metabolism^101–106^. These metabolic effects of *APOE4* are mediated by sex, with a stronger metabolic perturbation in *APOE4* females^105,106^. Like APOE, TREM2 signaling regulates metabolism. TREM2 loss of function results in low ATP, glycolytic deficits, and increased autophagy in microglia^107,108^. While the effects of *R47H* on metabolism are not well characterized, one study has found that *R47H* reduces both mitochondrial and glycolytic metabolic ability in microglia and impairs the immunometabolic switch to glycolysis upon inflammation^109^. Of note, both *APOE4* and *R47H* risk variants have been linked to increases in oxidative stress in human AD patients^110–113^. Reactive oxygen species (ROS) production can trigger mtDNA release into the cytosol and render it more resistant to degradation^114–116^. Since *APOE4* and *R47H* are associated with increased oxidative stress^110–113^, the combination of *APOE4* and *R47H* may exacerbate ROS and mtDNA release upstream of cGAS-STING activation in our model. Taken together, we posit that *APOE4-R47H* alters mitochondrial membrane permeabilization, resulting in increased BAX-mediated mtDNA leakage and DNA damage, thereby stimulating the cGAS-STING pathway and associated senescence.

In addition to mtDNA release, the cGAS-STING pathway can be triggered by the recognition of cytosolic chromatin fragments (CCFs) released during DNA damage and early-stage senescence^73,74^. Degradation of the nuclear envelope lamina protein Lamin-B1 during senescence facilitates CCF release^117,118^, stimulating the cGAS-STING pathway and resulting in SASP production^72–74^. APOE can localize to the nucleus to bind DNA, directly affecting gene transcription^119,120^. A recent study found that APOE accumulates in senescent cells and drives the senescence phenotype by binding nuclear lamina proteins and disrupting nuclear architecture and heterochromatin organization^121^. It is possible this APOE-mediated nuclear envelope disruption also elevates CCF release and induces cGAS signaling. A study has found increased nuclear localization of APOE4 compared to APOE3^120^, indicating that APOE4 may alter levels of APOE-mediated DNA damage and senescence. Future studies should dissect the relationship between APOE and senescence and determine if the *APOE4* risk allele alters this pathway.

The exact mechanisms underlying *APOE4*- and *R47H*-conferred AD risk remain elusive, despite their recognition as two of the strongest AD risk factors. Our study highlights interactions between *APOE4* and *R47H* risk alleles in microglia, such that their combination stimulates cell-autonomous cGAS-STING signaling and IFN-I response, resulting in cGAS-dependent senescence. While the mechanistic underpinnings of *APOE4-R47H*-mediated induction of cGAS-STING signaling and senescence remain murky, the efficacy of BAX inhibition at ameliorating senescence, along the known roles of APOE and TREM2 in mitochondrial health, point to a crucial role for BAX-mediated mitochondrial membrane permeabilization in this effect. A major limitation of our study is the lack of human AD patient data from *APOE4* and *R47H* female carriers. A previous study has found that in cases of sporadic AD, *R47H* carriers upregulate IFN-I response and significantly increase senescent microglia in the hippocampus, but this patient data was unfortunately not stratified by *APOE* allele^30^. Taken together, cGAS-STING and senescence pathways are known drivers of neurodegeneration^63,67,68,71,95^, and our study further links upregulation of these pathways to the strongest AD risk factors. This evidence points to cGAS-STING and senescence as vital pathological mechanisms in AD and important targets for future therapeutics.

## ACKNOWLEDGEMENTS

This work was supported by the NIH (F31AG079560 to G.C., R01AG076448 to L.G., R01AG072758 to L.G., R01AG054214 to L.G., R01AG074541 to L.G. and S.C.S., R01AG064239 to W.L., and K99AG078493 to S.A.), Tau Consortium and JPB Foundation (to L.G.), CureAD fund (to L.G. and S.C.S.), Daedalus fund (to L.G. and S.C.S.), BrightFocus Foundation Postdoctoral Fellowship in Alzheimer’s Disease Research (A20201312F to S.A.), and JumpStart Research Career Development Program (to S.A.).

## AUTHOR CONTRIBUTIONS

L.G. and G.C. conceived the project and planned experiments. L.G., G.C., W.L., L.F., and S.A. designed experiments. G.C., L.F., N.F., K.N., P.Y., M.Y.W., D.Z., F.Y., J.X., and F.C. performed experiments or analyses. S.C.S, A.Y., S.M., X.C., D.M.H., E.Z., and E.G. developed experimental protocols, tools, or reagents. G.C., M.Y.W., H.C., Y.H., S.A., and E.Z. provided technical support and troubleshooting. G.C., K.N., P.Y., and M.Y.W. maintained the mouse colony. S.G. designed and created the *hTREM2-CV* and *hTREM2-R47H* knock-in mouse lines. G.C. and L.G. wrote the manuscript. All authors read and approved the paper.

## STAR ★ METHODS

### KEY RESOURCES

| REAGENT OR RESOURCE | SOURCE | IDENTIFIER |
| --- | --- | --- |
| **Mice** |  |  |
| Prnp-MAPT*P301S (B6/C3) | The Jackson Laboratory | RRID:IMSR_JAX:008169 |
| C57BL/6J | The Jackson Laboratory | RRID:IMSR_JAX:000664 |
| Human APOE3 or APOE4 (C57BL/6J) | P.M. Sullivan, Duke University | Sullivan et al., 1997 (PMID: 9218423) |
| Human TREM2-CV (C57BL/6J) | Gan Lab, Weill Cornell; this paper | N/A |
| Human TREM2-R47H (C57BL/6J) | Gan Lab, Weill Cornell; Sayed et al., 2021 | Sayed et al., 2021 (PMID: 34851693) |
| **Antibodies** |  |  |
| Rabbit anti-VGAT | Millipore Sigma | Cat. # ab5062P; RRID:AB_2301998 |
| Mouse anti-MC1 Tau | Dr. Peter Davies, Albert Einstein College of Medicine | Cat. # MC1; RRID: AB_2314773 |
| Rabbit anti-STING | Cell Signaling Technology | Cat. # 13647; RRID:AB_2732796 |
| Rabbit anti-phospho-Stat1 (Tyr701) | Cell Signaling Technology | Cat. # 9167; RRID:AB_561284 |
| Goat anti-Iba1 | Abcam | Cat. # ab5076; RRID:AB_2224402 |
| Rabbit anti-p21 (used on mouse brain sections) | Abcam | Cat. # ab188224; RRID:AB_2734729 |
| Rabbit anti-p21 (used on primary cells) | Abcam | Cat. # ab109199; RRID:AB_10861551 |
| Mouse anti-γH2AX, (phospho S139) | Abcam | Cat. # ab26350; RRID:AB_470861 |
| Rabbit anti-cGAS (D3O8O) | Cell Signaling Technology | Cat. # 31659; RRID:AB_2799008 |
| Mouse anti-GAPDH | GeneTex | Cat. # GTX627408; RRID:AB_11174761 |
| Mouse anti-β-actin | Sigma-Aldrich | Cat. # A3854; RRID:AB_262011 |
| Peroxidase-AffiniPure Goat anti-rabbit IgG | Jackson ImmunoResearch Labs | Cat. ## 111-035-144; RRID:AB_2307391 |
| Alexa Fluor 488 Donkey Anti-Goat IgG | Thermo Fisher Scientific | Cat. # A-11055; RRID:AB_2534102 |
| Alexa Fluor 488 Donkey Anti-Rabbit IgG | Thermo Fisher Scientific | Cat. # A-21206; RRID:AB_2535792 |
| Alexa Fluor 568 Donkey Anti-Mouse IgG | Thermo Fisher Scientific | Cat. # A10037; RRID:AB_11180865 |
| Alexa Fluor 568 Donkey Anti-Rabbit IgG | Thermo Fisher Scientific | Cat. # A10042; RRID:AB_2534017 |
| Alexa Fluor 488 Goat Anti-Rabbit IgG | Thermo Fisher Scientific | Cat. # A-11008; RRID:AB_143165 |
| Alexa Fluor 568 Goat Anti-Mouse IgG | Thermo Fisher Scientific | Cat. # A-11031; RRID:AB_144696 |
| **Chemicals, peptides, and recombinant proteins** |  |  |
| Cas9 protein | Integrated DNA technologies | Cat. # 1081058 |
| β-mercaptoethanol | Sigma-Aldrich | Cat. # M3148 |
| RNase-free DNase | Qiagen | Cat. # 79254 |
| SuperScript IV VILO Master Mix | Invitrogen | Cat. # 11756050 |
| SsoAdvanced Universal SYBR Green Supermix | BioRad | Cat. # 1725270 |
| Bovine serum albumin | Sigma-Aldrich | Cat. # A3294 |
| DL-1,4-Dithiothreitol (DTT) | Thermo Fisher Scientific | Cat. # 426380500 |
| Ambion RNase inhibitor | Invitrogen | Cat. # AM2682 |
| Paraformaldehyde | Electron Microscopy Services | Cat. # 15710-S |
| PBS | Gibco | Cat. # 70011-044 |
| Triton X-100 | Sigma-Aldrich | Cat. # T9284 |
| TBST | Thermo Fisher Scientific | Cat. # 28360 |
| VectaShield antifade mounting medium with DAPI | Vector Laboratories | Cat. # H-1200 |
| VectaMount AQ aqueous mounting medium | Vector Laboratories | Cat. # H-5501 |
| Donkey serum | Jackson ImmunoResearch | Cat. # 017-000-121 |
| Goat serum | Jackson ImmunoResearch | Cat. # 005-000-121 |
| Sudan black B, fat soluble diazo dye | Abcam | Cat. # ab146284 |
| Trypsin | Gibco | Cat. # 15090-45 |
| DNase I | Millipore Sigma | Cat. # 260913 |
| DMEM | Corning | Cat. # 10-017-CV |
| Fetal bovine serum | Gibco | Cat. # 26140-079 |
| Penicillin-streptomycin | Gibco | Cat. # 15070063 |
| Poly-D-lysine | Thermo Fisher Scientific | Cat. # A389401 |
| Recombinant mouse GM-CSF, carrier free | R&D Systems | Cat. # 415-ML-020-CF |
| 0N4R tau fibrils | Dr. Sue-Ann Mok, University of Alberta | N/A |
| RIPA lysis and extraction buffer | Thermo Fisher Scientific | Cat. # 89901 |
| Halt phosphatase inhibitor cocktail | Thermo Fisher Scientific | Cat. # 78420 |
| cOmplete protease inhibitor cocktail | Roche | Cat. # 11697498001 |
| NuPAGE MES SDS running buffer | Invitrogen | Cat. # NP0002 |
| Clarity Western ECL substrate | Bio-Rad | Cat. # 1705060 |
| Calcein AM | Invitrogen | Cat. # C3099 |
| Etoposide | Sigma-Aldrich | E1383 |
| TDI6570 (cGAS inhibitor) | Synthesized by Gan Lab at Wuxi AppTec (Tianjin) Co., Ltd. | Lama et al., 2019 (PMID: 31113940) |
| BAI1 (BAX channel blocker) | Sigma-Aldrich | Cat. # 196805; Garner et al., 2019 (PMID: 30718816) |
| Dimethyl sulfoxide (DMSO) | MP Biomedicals | Cat. # 02196055 |

| **Critical commercial assays** |  |  |
| --- | --- | --- |
| PureLink RNA Mini Kit | Invitrogen | Cat. # 12183018A |
| Simple Plex NFL Assay Kit | BioTechne | Cat. # SPCKB-PS-002448 |
| Nuclei Isolation Kit: Nuclei PURE Prep | Sigma-Aldrich | Cat. # NUC201-1KT |
| Chromium Single Cell 3' Reagent Kit | 10x Genomics | Cat. # PN-1000268 |
| MOM Basic Immunodetection Kit | Vector Laboratories | Cat. # BMK-2202 |
| Quick-RNA MicroPrep Kit | Zymo Research | Cat. # R1051 |
| BCA protein assay kit | Pierce | Cat. # 23225 |
| Multiplex mouse cytokine/chemokine 32-plex kit | EMD Millipore | Cat. # MCYTMAG-70K-PX32 |
| BrdU cell proliferation ELISA kit | Abcam | Cat. # ab126556 |
| **Deposited data** |  |  |
| Immunoblot data | This paper | N/A |
| Raw and processed data (bulk and single-nuclei RNA-sequencing) | This paper | N/A |
| **Software and algorithms** |  |  |
| R 4.2.3 | The R project | https://www.r-project.org/ |
| RStudio 2023.06.0+421 | RStudio | https://rstudio.com |
| Gene set enrichment analysis (GSEA) | Subramanian et al., 2005; PMID: 16199517 | https://www.gsea-msigdb.org/gsea/msigdb/index.jsp |
| Ingenuity pathway analysis | Qiagen | https://digitalinsights.qiagen.com/products-overview/discovery-insights-portfolio/analysis-and-visualization/qiagen-ipa/ |
| Cell Ranger 6.1.2 | 10x Genomics | https://support.10xgenomics.com/single-cell-gene-expression/software/pipelines/latest/what-is-cell-ranger |
| Seurat 4.0.0 | Seurat | https://satijalab.org/seurat/ |
| DESeq2 1.38.3 | Love et al., 2014; PMID: 25516281 | https://bioconductor.org/packages/release/bioc/html/DESeq2.html |
| Fiji/ImageJ 2.14.0 | Fiji | RRID: SCR_002285 |
| Image Lab 6.0.0 | Bio-Rad | https://www.bio-rad.com/en-us/product/image-lab-software?ID=KRE6P5E8Z |
| Adobe Illustrator 28.0 | Adobe | https://www.adobe.com/products/illustrator.html |
| GraphPad Prism 10 | Prism v10.1.0 | https://www.graphpad.com |
| Zen 3.2 (blue edition) | Zeiss | https://www.zeiss.com/microscopy/en/products/software/zeiss-zen.html |

### Mice

Human *TREM2-R47H* mice were generated using CRISPR/Cas9, as described previously^29^. *R47H/+* mice were crossed with human *APOE3/APOE3* or *APOE4/APOE4* knock-in mice (C57BL/6 background, provided by P.M. Sullivan, Duke University)^122^ to generate *APOE4/mApoe R47H/+* or *APOE3/mApoe R47H/+* mice. *APOE3* and *APOE4* lines were maintained separately. Subsequent crossing of F1 litters generated *APOE4/APOE4 R47H/+* and *APOE3/APOE3 R47H/+* mice. These F2 mice were then crossed with *P301S* tau transgenic mice (B6/C3 background, The Jackson Laboratory, RRID:IMSR_JAX:008169) to generate *APOE4/mApoe R47H/+ P301S+* and *APOE3/mApoe R47H/+ P301S+* mice. F3 crossing resulted in *APOE4/APOE4 R47H/+ P301S+* or *APOE3/APOE3 R47H/+ P301S+* mice. Crossing of F4 litters for the APOE4 line generated four genotypes of littermate mice for experiments: *APOE4/APOE4 mTrem2+/+, APOE4/APOE4 mTrem2+/+ P301S+, APOE4/APOE4 R47H/+, APOE4/APOE4 R47H/+ P301S+*. Crossing of F4 litters for the APOE3 line generated four genotypes of littermate mice for experiments: *APOE3/APOE3 mTrem2+/+, APOE3/APOE3 mTrem2+/+ P301S+, APOE3/APOE3 R47H/+, APOE3/APOE3 R47H/+ P301S+*.

CRISPR/Cas9-mediated knock-in of human *TREM2* common variant (CV) cDNA into the mouse *Trem2* genetic locus was done by pronuclear injection. Briefly, a single guide RNA was selected and RNP containing sgRNA (Synthego) and Cas9 protein (Integrated DNA Technologies, 1081058) was prepared. A donor plasmid was designed and made (GenScript). The human *TREM2* cDNA and a poly A tail were flanked on each side by 0.8-kb homology arms that were obtained from the *mTrem2* genomic DNA sequence. Pronuclear injections of RNP and the donor plasmid were performed at the Rockefeller University transgenic core facility with embryos from C57BL/6J (The Jackson Laboratory, RRID:IMSR_JAX:000664). The *Trem2* target region was CTGATCACAGGTGGGACCAC and the sgRNA sequence (sense) was CTGATCACAGGTGGGACCAC. Genomic DNA from founders was isolated from tail lysate. To screen the specific knock-in mice, two PCR amplifications were performed with primers (Integrated DNA Technologies) flanking the outside of the homology arms and on the human *TREM2* transgene. The PCR products were Sanger sequenced (Azenta) to validate the correct insertion and the locus integrity. Specific integration of the donor DNA was characterized further, and no non-specific integration was detected.

To distinguish homozygous from heterozygous mice, tail DNA was characterized by PCR with three primers, two primers on the targeting genomic sequences and one primer on the transgene. The *hTREM2- CV* line was maintained independently and backcrossed to non-transgenic C57BL/6 mice for two to three generations, then crossed with *APOE4/APOE4* KI mice to generate *APOE4/mApoe CV/+* mice. Subsequent crossing of F1 litters generated *APOE4/APOE4 CV/CV* mice. These F2 mice were crossed with the *APOE4/APOE4 R47H/+* mice described previously to generate *APOE4/APOE4 R47H/CV* mice. F3 crossing resulted in *APOE4/APOE4 R47H/CV* and *APOE4/APOE4 CV/CV* mice, which were used for breeding to produce *APOE4/APOE4 R47H/CV* and *APOE4/APOE4 CV/CV* littermate mice for experiments.

Mice were housed with no more than five animals per cage, given ad libitum access to food and water, and housed in a pathogen-free barrier facility at 21–23 °C with 30–70% humidity on a 12-hour light/12-hour dark cycle. At 9–10 months of age, mice were used for pathology and RNA-seq studies. Mice of both sexes were used for pathology, and analyses based on sex are included in the supplementary figures. Female mice were used for snRNA-Seq studies. All mouse protocols were approved by the Institutional Animal Care and Use Committee at Weill Cornell Medicine.

### Quantitative reverse-transcription PCR

Snap-frozen frontal cortices were thawed and homogenized with a 27G needle in lysis buffer provided in the PureLink RNA Mini Kit (Invitrogen, 12183018A) with 1% β-mercaptoethanol (Sigma, M3148). RNA was isolated with the PureLink RNA Mini Kit, and the remaining DNA was removed by incubation with RNase-free DNase (Qiagen, 79254). Purified mRNA was then converted to cDNA with the SuperScript IV VILO Master Mix (Invitrogen, 11756050). Quantitative RT-PCR was performed on the CFX96 real-time PCR detection system (BioRad) with SsoAdvanced Universal SYBR Green Supermix (BioRad, 1725270). The average value of three replicates for each sample was expressed as a threshold cycle (Cq), the point at which the fluorescent signal starts to increase rapidly. Then, the difference (ΔCtq) between Cq values for the transcript of interest and for mouse Gapdh was calculated for each sample. The gene expression for each sample was calculated using the 2–ΔΔCq method and reported as fold change relative to the average of the mouse *Trem2+/+* sample. The following primers were used: Primer: Human TREM2 Fwd: CACTCTCACCATTACGCTGC Primer: Human TREM2 Rev: GGAACCAGAGATCTCCAGCA Primer: Mouse TREM2 Fwd: ACCGTCACCATCACTCTGAA Primer: Mouse TREM2 Rev: GGTGGGAAGGAGGTCTCTTG Primer: Mouse GAPDH Fwd: GGAGCGAGACCCCACTAACA Primer: Mouse GAPDH Rev: ACATACTCAGCACCGGCCTC

### Plasma neurofilament light measurement

Blood was collected from mice via cardiac puncture just prior to perfusion. Blood was centrifuged at 5000 rpm for 5 minutes, and the plasma supernatant was collected and snap-frozen on dry ice. Plasma was diluted 1:5, and NFL concentration was measured using the Ella-SimplePlex^TM^ NFL assay kit (BioTechne, SPCKB-PS-002448)^123^.

### Nuclei isolation from frozen mouse hippocampi

Nuclei isolation from frozen mouse hippocampi from transcardially perfused mice was adapted from a previous study, with modifications^124,125^. All procedures were performed on ice or at 4 °C. In brief, postmortem mouse hippocampal tissue was placed in 1,500 μl of nuclei PURE lysis buffer from the nuclei PURE prep kit (Sigma, NUC201) and homogenized with a Dounce tissue grinder (Sigma, D8938-1SET) with 15 strokes with pestle A and 15 strokes with pestle B. The homogenized tissue was filtered through a 35-μm cell strainer, centrifuged at 600g for 5 min at 4 °C and washed three times with 1 ml of PBS containing 1% bovine serum albumin (BSA; Sigma, A7906), 20 mM DTT (Thermo Fisher Scientific, 426380500), and 0.2 U μl^−1^ recombinant Ambion RNase inhibitor (Invitrogen, AM2682). Nuclei were then centrifuged at 600g for 5 min at 4 °C and resuspended in 500 μl of PBS containing 0.04% BSA and 1x DAPI, followed by fluorescence-activated cell sorting to remove cell debris. The sorted suspension of DAPI-stained nuclei was counted and diluted to a concentration of 1,000 nuclei per μl in PBS containing 0.04% BSA.

### Droplet-based snRNA-seq

For droplet-based snRNA-seq, libraries were prepared with Chromium Single Cell 3′ Reagent kits (v3.1; 10x Genomics, PN-1000268), according to the manufacturer’s protocol. The snRNA-seq libraries were sequenced on a NovaSeq 6000 sequencer (Illumina) with PE 2 x 50 paired-end kits by using the following read length: 28 cycles Read 1, 10 cycles i7 index, 10 cycles i5 index, and 90 cycles Read 2.

### Analysis of droplet-based snRNA-seq data from hippocampal tissue

We sequenced and integrated samples from *E4/E4/+/+* (n = 3), *E4/E4/+/+/P301S* (n = 3), *E4/E4/R47H/+/P301S* (n = 3), *E3/E3/+/+* (n = 3), *E3/E3/+/+/P301S* (n = 3), and *E3/E3/R47H/+/P301S* (n = 3) mice. Gene counts were obtained by aligning reads to the mm10 genome with Cell Ranger software (v6.1.2; 10x Genomics). To account for unspliced nuclear transcripts, reads mapping to pre-mRNA were counted. Cell Ranger 6.1.2 default parameters were used to call cell barcodes. We further removed genes expressed in no more than 3 cells, cells with unique gene counts over 5,000 or less than 300, cells with UMI counts over 15,000 and cells with a high fraction of mitochondrial reads (>1%). Potential doublet cells were predicted using DoubletFinder^57^ for each sample separately, with high-confidence doublets removed. Normalization and clustering were done with the Seurat package v4.0.0. In brief, counts for all nuclei were scaled by the total library size multiplied by a scale factor (10,000) and transformed to log space. A set of 2,000 highly variable genes was identified with SCTransform from the sctransform R package in the variable stabilization mode. This returned a corrected unique molecular identifier count matrix, a log-transformed data matrix and Pearson residuals from the regularized negative binomial regression model. Principal-component analysis was done on all genes, and t-distributed stochastic neighbor embedding was run on the top 15 principal components. Cell clusters were identified with the Seurat functions FindNeighbors (using the top 15 principal components) and FindClusters (resolution = 0.1). In this analysis, the neighborhood size parameter pK was estimated using the mean variance-normalized bimodality coefficient (BCmvn) approach, with 15 principal components used and pN set as 0.25 by default. For each cluster, we assigned a cell-type label using statistical enrichment for sets of marker genes and manual evaluation of gene expression for small sets of known marker genes^126,127^. Differential gene expression analysis was done using the FindMarkers function and MAST^128^. To identify gene ontology and pathways enriched in the DEGs, DEGs were analyzed using the MSigDB gene annotation database^129,130^. To identify gene activation networks and upstream regulators, DEGs were analyzed using Ingenuity Pathway Analysis (QIAGEN, Inc.). To control for multiple testing, we used the Benjamini–Hochberg approach to constrain the FDR.

### Immunohistochemistry and image analysis

Hemibrains from transcardially perfused mice were placed in 4% paraformaldehyde (Electron Microscopy Services, 15710-S) for 48-hours, followed by 30% sucrose in PBS for 48 hours at 4°C. Sections were cut coronally at 30 μm using a freezing microtome (Leica SM2010) to generate eight free-floating series per mouse and placed in cryoprotective medium at –20°C until use. All washing steps were repeated three times for 5 minutes each. For VGAT (Millipore Cat# AB5062P, RRID:AB_2301998, 1:400 dilution), MC1 (Custom, RRID: AB_2314773, 1:2500 dilution), and STING (Cell Signaling Technology Cat# 13647, RRID:AB_2732796, 1:200 dilution) antibodies, sections were washed in PBS, permeabilized with TBST (0.5% Triton X-100; Thermo Fisher Scientific, 28360) for 10 minutes, washed again, and immersed in 10% BSA (Sigma, A7906) in TBST for 30 minutes. A mouse-on-mouse blocking step was performed using 2% MOM Blocking Reagent (MOM Basic Immunodetection Kit, Vector Laboratories, BMK-2202) in PBS for 1 hour at room temperature. Samples were then washed and incubated with 8% MOM Protein Concentrate in PBS for 5 minutes. Primary antibodies were diluted in 8% MOM Protein Concentrate in PBS and incubated overnight at 4°C. The following day, sections were washed and incubated in appropriate secondary antibodies diluted in 8% MOM Protein Concentrate in PBS for 1 hour. Sections were washed, mounted on slides using Vectashield antifade mounting medium with DAPI (Vector Laboratories, H-1200), and imaged.

For p-STAT1 (Cell Signaling Technology Cat# 9167, RRID:AB_561284, 1:400 dilution), IBA1 (Abcam Cat# ab5076, RRID:AB_2224402, 1:300 dilution), and p21 (Abcam Cat# ab188224, RRID:AB_2734729, 1:100 dilution) antibodies, sections were washed in PBS, permeabilized with TBST (0.5% Triton X-100) for 10 minutes, washed again, and immersed in 5% Donkey Serum (Jackson ImmunoResearch, 017-000-121) in TBST for 2 hours at room temperature. Sections were washed, followed by incubation with primary antibodies diluted in 5% Donkey Serum in TBST overnight at 4°C. The following day, sections were washed and incubated in appropriate secondary antibodies in 1% BSA in TBST for 1 hour. Secondary antibodies used were the following (1:500 dilution): Alexa Fluor 488 Donkey Anti-Goat IgG (Thermo Fisher Scientific Cat# A-11055, RRID:AB_2534102), Alexa Fluor 488 Donkey Anti-Rabbit IgG (Thermo Fisher Scientific Cat# A-21206, RRID:AB_2535792), Alexa Fluor 568 Donkey Anti-Mouse IgG (Thermo Fisher Scientific Cat# A10037, RRID:AB_11180865), Alexa Fluor 568 Donkey Anti-Rabbit IgG (Thermo Fisher Scientific Cat# A10042, RRID:AB_2534017). Sections were washed, mounted on slides using Vectashield antifade mounting medium with DAPI (Vector Laboratories, H-1200), and imaged.

For VGAT, the CA1 region of the mouse hippocampus was imaged with a LSM880 confocal microscope and Zen black image acquisition software using a ×40 objective. Sections were imaged with a 1-μm interval z-stack over a total distance of 6 μm per slice with XY stitching. For MC1 and IBA1, the hippocampus was imaged with a Keyence BZ-X700 microscope using a ×20 objective with XY stitching. For pSTAT1, STING, and p21, the CA1 or CA3 region of the mouse hippocampus was imaged with a Zeiss Apotome microscope using a ×20 objective. Sections were imaged with a 1-μm interval z-stack over a total distance of 8 μm per slice. Final images were processed with maximum intensity projection. Quantification was done using ImageJ software (NIH).

### Brain volumetric analysis

Brain volumetric quantification was adapted from a previous study, with modifications^39^. Hippocampal sections spanning from bregma ∼1.3 mm to bregma ∼3.1 mm were selected and mounted from one series of brain sections (approximately 10-12 sections per animal with 240 μm between sections). Mounted sections were washed in Milli-Q water for 1 minute. Sections were then stained with 0.1% Sudan black (Abcam, ab146284) in 70% ethanol at room temperature for 30 minutes, washed in 70% ethanol three times for 10 seconds each, washed in Milli-Q water three times for 2 minutes each, and were coverslipped with VectaMount mounting media (Vector laboratories, H-5501). Slides were imaged with a Keyence BZ- X700 microscope using a ×4 objective on brightfield with XY stitching. ImageJ software was used to trace regions of interest and quantify area. Volume within each animal for the region of interest was calculated using the following formula: volume = (sum of area) x 240 μm.

### Isolation and culture of primary microglia

Cortices were dissected from mouse pups at postnatal day 1-3. The meninges were removed, and the cortical tissue was finely chopped with a razor blade and digested in 0.1% trypsin (Gibco, 15090-45) with DNase I (Millipore Sigma, 260913) at 37°C for 22 minutes. Digestion was stopped with DMEM (Corning, 10-017-CV) containing 20% heat inactivated FBS (Gibco, 26140079). 200 μl of DNAse was added, and tissue was triturated and spun at 200 g for 15 minutes. The pellet was resuspended in DMEM supplemented with 10% heat inactivated FBS (Gibco, 26140-079) and 1% penicillin-streptomycin (Gibco, 15070063) and was plated in T75 flasks precoated with 5 μg/ml poly-D-lysine (Thermo Fisher Scientific, A389401) and rinsed with water. Mixed glial cultures were maintained in growth media (DMEM with 10% FBS and 1% P/S) at 37°C and 5% CO2 for 8-10 days. Once bright, round cells began to appear in the culture on day 8-10, media was supplemented with 5 ng/ml recombinant mouse GM-CSF (R&D Systems, 415-ML-020-CF) to promote microglial proliferation. When flasks were ready for harvest on day 12-14, to isolate microglia, the flasks were shaken at 400 rpm for 2 hours, media was spun down at 200 g for 15 minutes, and cells were plated for experiments on poly-D-lysine coated plates.

### Bulk RNA-Seq and analysis of primary microglia

Primary microglia were plated at a density of 400,000 cells/well in a PDL-coated 24-well plate in growth media. The next day, cells were treated with 1 μg/ml of 0N4R tau fibrils (from Dr. Sue-Ann Mok, University of Alberta) per well for 18 hours. RNA was isolated from microglia using the Quick-RNA MicroPrep Kit (Zymo Research, R1051). RNA was shipped to Novogene for library preparation and bulk RNA sequencing. Differential gene expression was analyzed with the DESeq2 1.38.3 package^131^. Counts were normalized using the median of ratios method. Genes with <15 counts across all samples were excluded from analysis. To control for multiple testing, we used the Benjamini–Hochberg approach to constrain the FDR. Pathway analysis was done using the MSigDB gene annotation database. Ingenuity Pathway Analysis (QIAGEN, Inc.) was used to identify gene activation networks and upstream regulators.

### Western blots

For mouse brain samples, hippocampal or frontal cortex tissue was mechanically homogenized on ice in RIPA buffer (Thermo Fisher Scientific, 89901) supplemented with Halt phosphatase inhibitor cocktail (Thermo Fisher Scientific, 78420) and cOmplete protease inhibitor cocktail (Roche, 11697498001). After sonication, brain lysates were centrifuged at 20,000 g for 15 minutes at 4°C. Supernatants were collected, and protein concentration was measured by BCA (Pierce, 23225).

For cultured primary microglia, cells were plated at a density of 500,000 cells/well in a PDL-coated 6-well plate in growth media. The next day, cells were treated with 1 μg/ml of 0N4R tau fibrils (from Dr. Sue-Ann Mok, University of Alberta) per well for 24 hours. Cells were lysed in RIPA buffer supplemented with Halt phosphatase inhibitor cocktail and cOmplete protease inhibitor cocktail. Samples were rotated for 15 minutes at 4°C, and lysates were cleared by centrifugation at 20,000 g for 10 minutes at 4°C. Protein concentration was measured by BCA.

50 μg of brain lysates or 10 μg of primary microglial lysates were loaded onto NuPAGE Bis-Tris gels (Invitrogen) and run in MES SDS running buffer (Invitrogen, NP0002) at 130 volts for approximately 1 hour. Gels were transferred to nitrocellulose membranes (Bio-Rad, 1620115) at 0.25 amps for 2 hours at 4°C. Membranes were washed three times for 10 minutes each in TBS with 0.01% Triton X-100 (TBST) and were blocked for 1 hour in 5% milk in TBST. Primary antibodies were diluted in 1% milk in TBST and incubated at 4°C overnight. The following day, membranes were washed three times for 10 minutes each with TBST and incubated with appropriate secondary antibodies in 1% milk in TBST for 1 hour at room temperature. Membranes were washed again three times for 10 minutes each in TBST to minimize non-specific binding. Membranes were then treated with ECL (Bio-Rad, 1705060) for 60 seconds and developed using a Bio-Rad imager. Blots were scanned at 300 d.p.i. and quantified using Image Lab (Bio-Rad). Primary antibodies used for western blot were anti-cGAS (Cell Signaling Technology Cat# 31659, RRID:AB_2799008, 1:500 dilution), anti-GAPDH (GeneTex Cat# GTX627408, RRID:AB_11174761, 1:10000 dilution), and anti-β-Actin (Sigma-Aldrich Cat# A3854, RRID:AB_262011, 1:10000 dilution).

Immunoreactivity was detected with goat anti-rabbit HRP (Jackson ImmunoResearch Labs Cat# 111-035- 144, RRID:AB_2307391, 1:5000 dilution).

### Cytokine secretion measurement

Primary microglia were plated at a density of 50,000 cells/well in a PDL-coated 96-well plate in growth media. The next day, cells were treated with 1 μg/ml of 0N4R tau fibrils (from Dr. Sue-Ann Mok, University of Alberta) per well for 18 hours. The conditioned media were collected and cleared by centrifugation at 3,000 g for 10 minutes at 4°C. The concentrations of cytokines/chemokines in the conditioned media were measured by a multiplex mouse cytokine/chemokine 32-plex kit (Millipore, MCYTMAG-70K-PX32) with the Luminex MAGPIX CCD Imager. For normalization, the live microglia were incubated in 2 μm Calcein AM (Invitrogen, C3099) for 20 minutes, and were imaged and counted with the SpectraMax i3X (Molecular Devices) using the MiniMax function. Cytokine secretion per well was divided by the cell count per well to normalize for cell density. The tau-induced fold change in cytokine secretion for each sample was calculated compared to the average cytokine secretion of unstimulated microglia.

### Microglial proliferation assay

Primary microglia were plated at a density of 50,000 cells/well in a PDL-coated 96-well plate in growth media. The next day, cells were treated with 1 μg/ml of 0N4R tau fibrils (from Dr. Sue-Ann Mok, University of Alberta) per well or 35 μm etoposide (Sigma, E1383) for 24 hours. Microglial proliferation was measured using a BrdU cell proliferation ELISA kit (Abcam, ab126556). BrdU label was added to each well at the time of treatment with tau or etoposide and was incubated for 24 hours. The level of BrdU incorporated per well was measured using the BioTek Synergy H1 plate reader.

### Immunocytochemistry and image analysis

Primary microglia were plated at a density of 75,000 cells/well in a PDL-coated chamber slide in growth media overnight. For γH2AX staining, cells were stimulated with 1 μg/ml of 0N4R tau fibrils (from Dr. Sue-Ann Mok, University of Alberta) per well for 72 hours. For p21 staining, cells were pre-treated with 20 μm cGAS inhibitor TDI6570^85^, 5 μm BAX inhibitor BAI1^84^, or 0.05% DMSO vehicle (MP Biomedicals, 02196055). After 4 hours of pre-treatment, cells were stimulated with 1 μg/ml of 0N4R tau fibrils (from Dr. Sue-Ann Mok, University of Alberta) per well for 72 hours. After the 72 hours of tau fibril treatment, for both γH2AX and p21 staining, cells were washed once in PBS and fixed in 4% paraformaldehyde for 15 minutes. Cells were washed three times with PBS and immersed in blocking solution consisting of 0.3% Triton-X- 100 (Sigma, T9284) and 5% goat serum (Jackson ImmunoResearch, 005-000-121) in PBS for 30 minutes. Cells were then incubated with primary antibody γH2AX (Abcam Cat# ab26350, RRID:AB_470861, 1:500 dilution) or p21 (Abcam Cat# ab109199, RRID:AB_10861551, 1:500 dilution) diluted in blocking solution overnight at 4°C. The following day, cells were washed three times with PBS, followed by incubation with secondary antibody Alexa Fluor 568 Goat Anti-Mouse (Thermo Fisher Scientific Cat# A-11031, RRID:AB_144696, 1:500 dilution) or Alexa Fluor 488 Goat Anti-Rabbit (Thermo Fisher Scientific Cat# A- 11008, RRID:AB_143165, 1:500 dilution) for 2 hours at room temperature. Cells were washed three times with PBS, wells were removed from the chamber slides, and the slides were cover-slipped in Vectashield antifade mounting medium with DAPI (Vector Laboratories, H-1200). Cells were imaged with a Keyence BZ-X700 microscope using a ×40 objective. Quantification was done using ImageJ software (NIH).

### Synthesis of cGAS inhibitor

TDI-6570 was prepared as described previously^85^. All intermediates were obtained by filtration using water and cold methanol. Purity of the products was confirmed by performing liquid chromatography– mass spectrometry analysis.

### Statistical Analysis

Statistical analyses were performed using GraphPad Prism v.10.1.0, and significant outliers were removed using Prism’s outlier calculator with Grubbs’ test (alpha = 0.05). All statistical details can be found in the figures and figure legends. A subset of mice from the immunohistochemistry cohort was randomly selected for snRNA-Seq studies. Experimenters were blinded of genotypes when performing image analysis and quantification for all immunohistochemistry and brain volumetric experiments.

## SUPPLEMENTARY FIGURES

**Figure S1.**
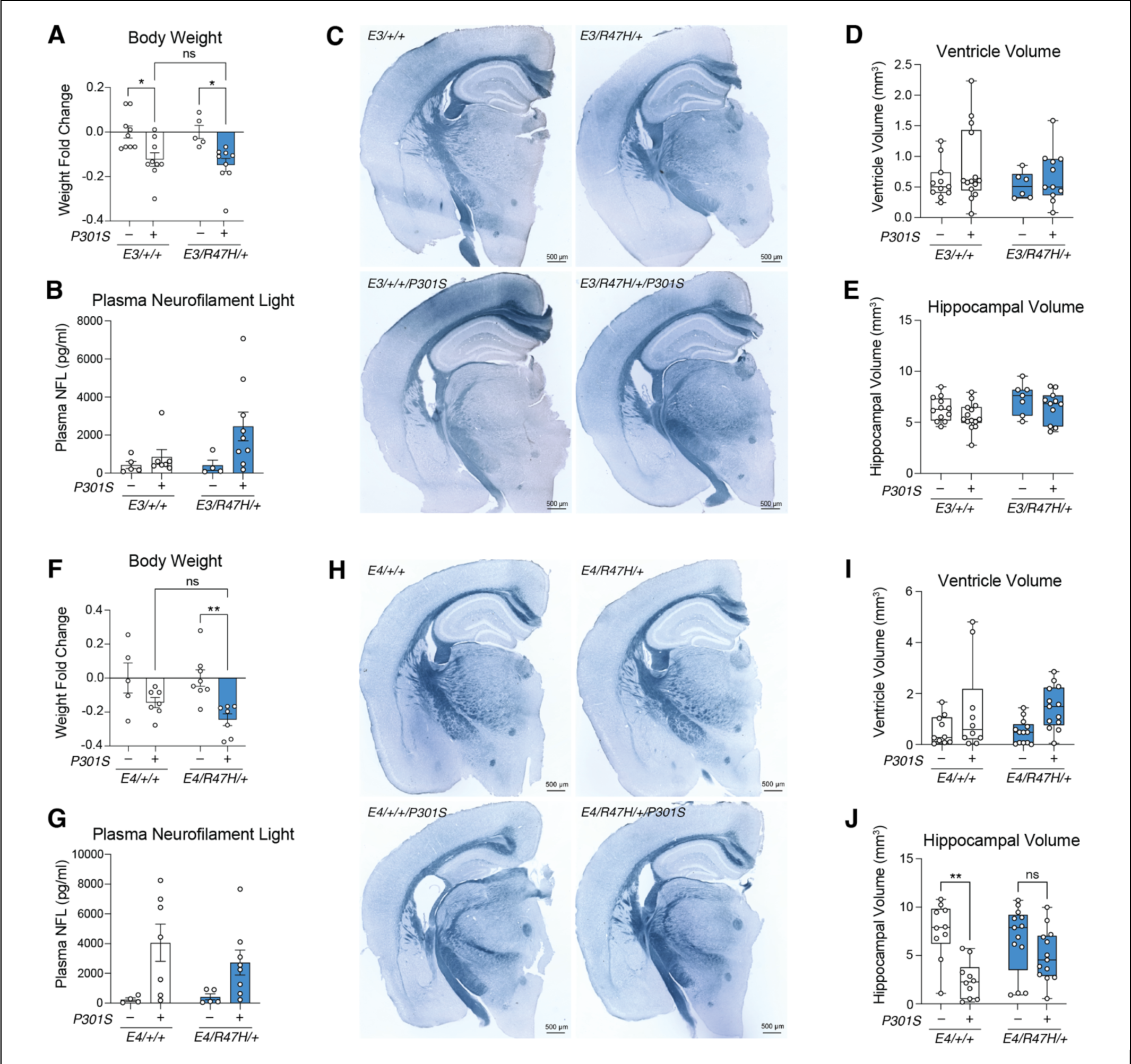
*APOE3-R47H* female mice and *APOE4-R47H* male mice do not worsen tauopathy-induced neurodegeneration at 9-months-old, related to Figure 1 (A) Quantification of body weight fold change in *P301S* transgenic versus non-transgenic female *APOE3/APOE3* mice within each genotype at 9-10 months old. Each circle represents one animal. * p < 0.05, n.s. not significant. Two-way ANOVA, Tukey’s multiple comparisons test. Data are reported as mean ± S.E.M. (B) Plasma neurofilament light concentration in female *APOE3/APOE3* mice. Each circle represents one animal. Two statistically significant outliers were removed. Two-way ANOVA, Tukey’s multiple comparisons test. Data not significant. Data are reported as mean ± S.E.M. (C) Representative images of female *APOE3/APOE3* mouse brain sections stained with Sudan black at 9-10 months old; scale bar, 500μm. (D and E) Quantification of ventricle volume (D) and hippocampal volume (E) in female *APOE3/APOE3* mice. Each circle represents sum volume measurements of eight to ten brain sections per animal. Two statistically significant outliers were removed for ventricle volume. Two-way ANOVA, Tukey’s multiple comparisons test. Data not significant. Data are reported as boxplot with min. to max. (F) Quantification of body weight fold change in *P301S* transgenic versus non-transgenic male *APOE4/APOE4* mice within each genotype at 9-10 months old. Each circle represents one animal. ** p = 0.0056, n.s. not significant. Two-way ANOVA, Tukey’s multiple comparisons test. Data are reported as mean ± S.E.M. (G) Plasma neurofilament light concentration in male *APOE4/APOE4* mice. Each circle represents one animal. One statistically significant outlier was removed. Two-way ANOVA, Tukey’s multiple comparisons test. Data not significant. Data are reported as mean ± S.E.M. (H) Representative images of male *APOE4/APOE4* mouse brain sections stained with Sudan black; scale bar, 500μm. (I and J) Quantification of ventricle volume (I) and hippocampal volume (J) in male *APOE4/APOE4* mice. Each circle represents sum volume measurements of nine to twelve brain sections per animal. ** p = 0.0014, n.s. not significant. Two-way ANOVA, Tukey’s multiple comparisons test. Data are reported as boxplot with min. to max.

**Figure S2.**
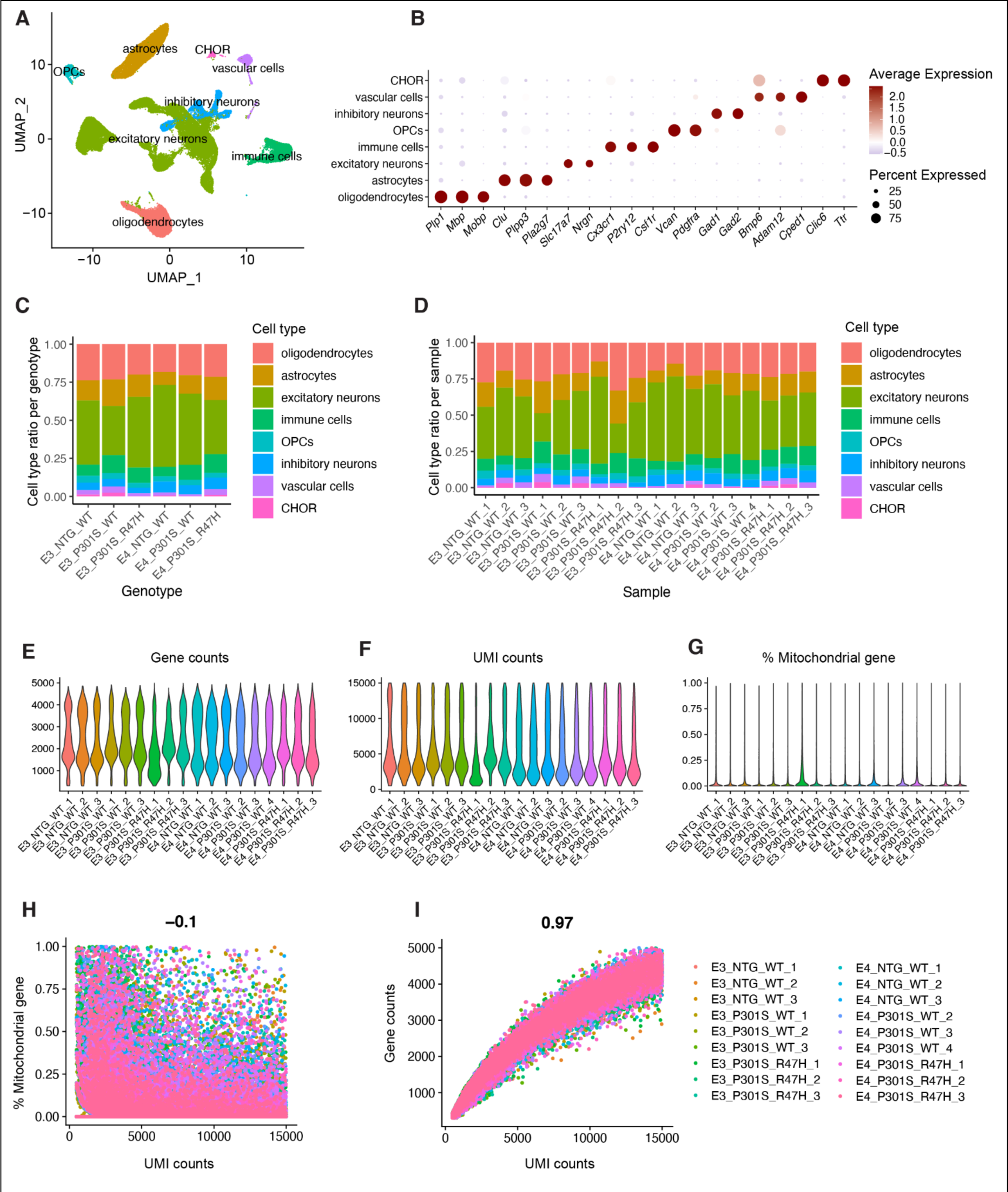
Quality control assessment of single-nuclei RNA-Seq, related to Figure 2 (A) UMAP plot of 91,744 single nuclei and their annotated cell types. OPC = oligodendrocyte precursor cells; CHOR = choroid plexus epithelial cells. (B) Dot plot showing expression of identity markers for each cell type. (C) Proportion of each cell type within differing *APOE* and *TREM2* genotypes. (D) Proportion of each cell type within individual samples. (E, F, and G) Violin plots showing total gene counts (E), UMI counts (F), and percentage of mitochondrial genes detected (G) per nuclei for each individual sample. (H and I) Correlation between UMI counts and percentage of mitochondrial genes detected (H) or total gene counts (I) per nuclei for each individual sample.

**Figure S3.**
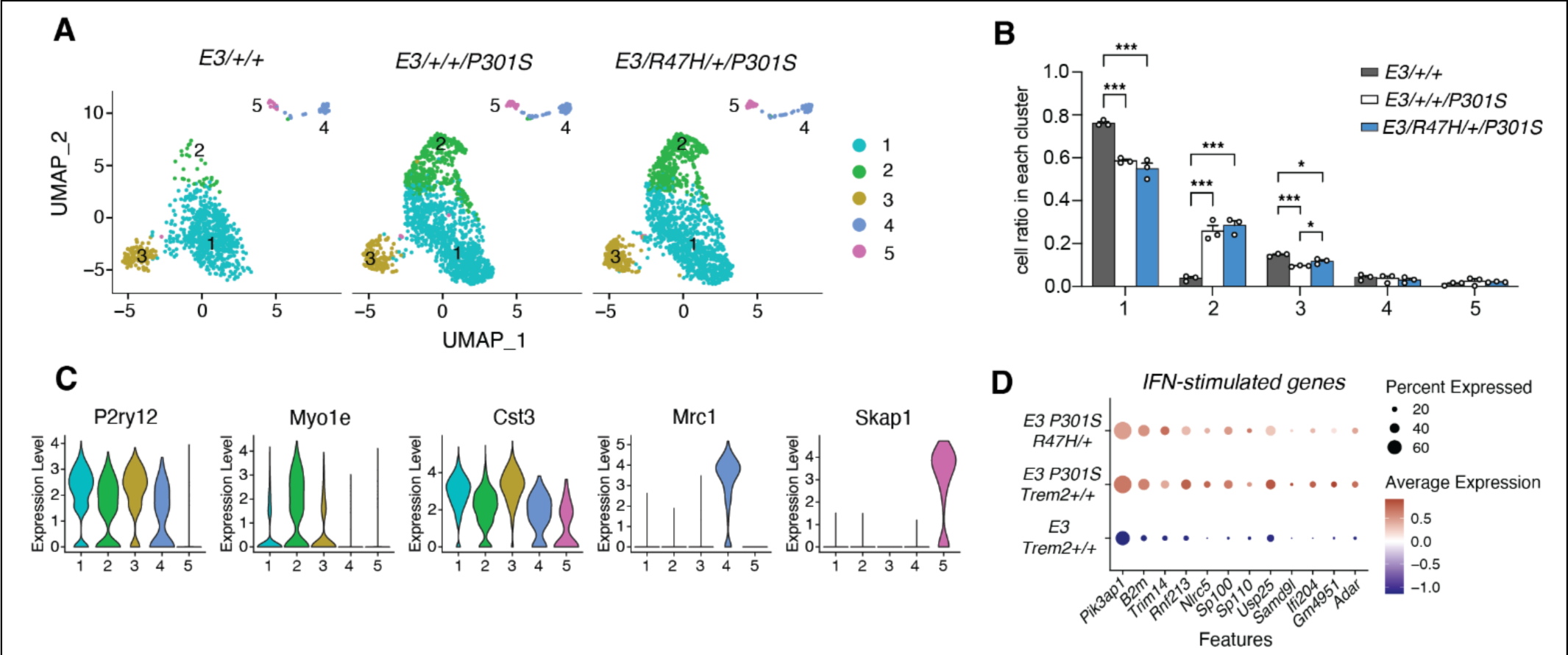
*APOE3*-*R47H* does not affect microglial interferon response, related to Figure 2 (A) UMAP plot of *APOE3* hippocampal immune cell nuclei split by genotype and colored according to subclusters. (B) Quantification of cell ratio per immune cell subcluster within each genotype. Each circle represents one animal. * p < 0.05, *** p < 0.001. One-way ANOVA, Tukey’s multiple comparisons test. Data are reported as mean ± S.E.M. (C) Violin plots showing cell type markers split by subcluster to highlight cluster identity. (D) Dot plot showing interferon-stimulated genes that were significantly enriched in *APOE4* microglial subcluster 4 and in *E4/R47H/+/P301S* versus *E4/+/+/P301S* hippocampal microglia. All genes listed are not significant in *E3/R47H/+/P301S* versus *E3/+/+/P301S* microglia.

**Figure S4.**
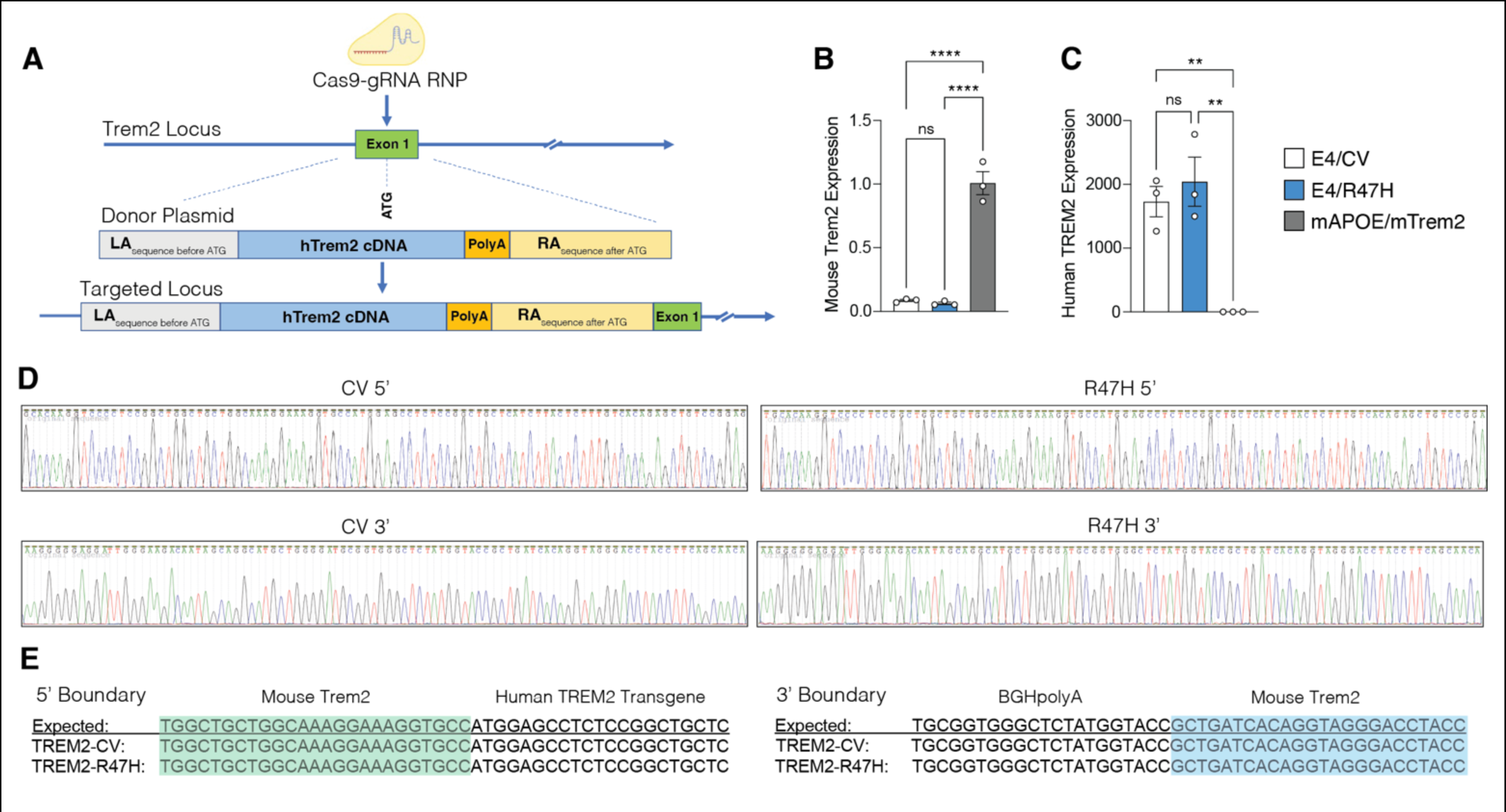
Characterization of *hTREM2-CV* and *hTREM2-R47H APOE4* mouse lines, related to Figure 5 (A) Schematic illustrating the CRISPR-mediated knock-in of human *TREM2-CV* and *TREM2-R47H* into C57BL/6J mouse *Trem2* genomic locus. (B and C) Bar plot showing PCR results for mouse *Trem2* expression (B) and human *TREM2* expression (C) in *APOE4-CV/CV*, *APOE4-R47H/CV*, or *mApoe/mTREM2* control mice. Each circle represents one animal. ** p < 0.01, **** p < 0.0001, n.s. not significant. One-way ANOVA, Tukey’s multiple comparisons test. Data are reported as mean ± S.E.M. (D) Sanger sequencing chromatogram for verifying the correct homozygous clone. Upper row shows 5’ end boundary, and lower row shows 3’ end boundary. (E) Sequences of the intended and obtained knock-in boundaries from PCR products with primers located outside of the homology arms and each end of the transgenes, showing correct homologous recombination. The 5’ end of the left homology arm is highlighted in green, and the 3’ end of the right homology arm is highlighted in blue.

**Figure S5.**
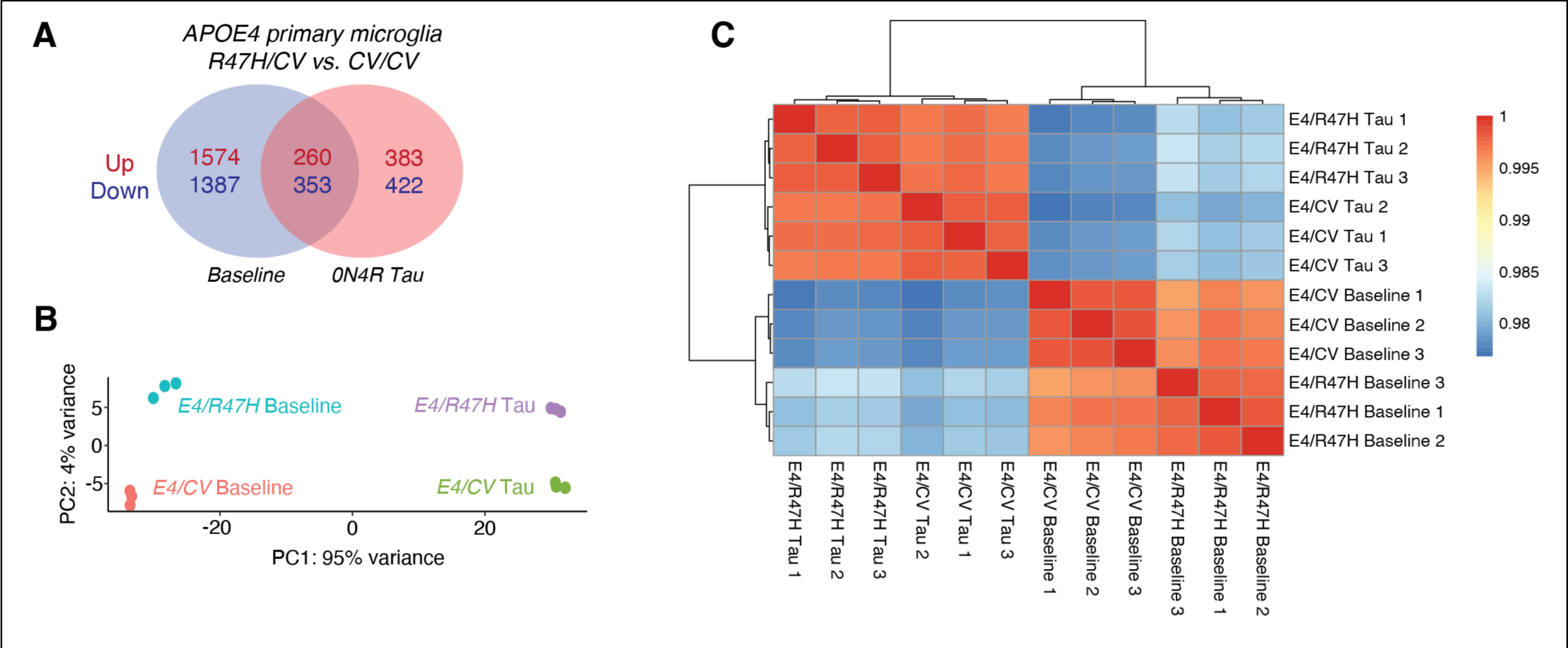
Quality control assessment of bulk RNA-Seq, related to Figure 5 (A) Venn diagram of *APOE4 R47H/CV* versus *CV/CV* primary microglia DEGs at baseline and after 18- hour 0N4R tau fibril stimulation (n = 3 biological replicates per condition). Red and blue numbers denote upregulated and downregulated DEGs, respectively (p < 0.05; log2foldchange β 0.1 or :: -0.1). (B) Principal component analysis plot. Tau fibril stimulation accounts for 95% of gene expression variance, and *TREM2* genotype accounts for 4% of gene expression variance. Each circle represents one biological replicate. (C) Heatmap showing correlations between each biological sample. Samples cluster based on *TREM2* genotype and tau stimulation status. There are no outlier samples.

## SUPPLEMENTARY INFORMATION

Supplementary Table 1 (S1)

snRNA-Seq markers for *APOE4* hippocampal immune cell subclusters. Bonferroni-adjusted *P* values were calculated using unpaired, two-tailed Wilcoxon rank-sum tests implemented in Seurat. Ingenuity pathway analysis upstream regulator predictions for microglial subcluster 4 (interferon) markers.

Supplementary Table 2 (S2)

DEGs in *APOE4-R47H-P301S* vs. *APOE4-WT-P301S* hippocampal mouse microglia. FDR-adjusted *P* values were calculated using MAST.

Supplementary Table 3 (S3)

snRNA-Seq markers for *APOE3* hippocampal immune cell subclusters. Bonferroni-adjusted *P* values were calculated using unpaired, two-tailed Wilcoxon rank-sum tests implemented in Seurat. DEGs in *APOE3- R47H-P301S* vs. *APOE3-WT-P301S* hippocampal mouse microglia. FDR-adjusted *P* values were calculated using MAST.

Supplementary Table 4 (S4)

DEGs in *APOE4-R47H-P301S* vs. *APOE4-WT-P301S* hippocampal mouse oligodendrocytes. FDR-adjusted *P* values were calculated using MAST. Gene set enrichment analysis hallmark pathways for *APOE4-R47H-P301S* vs. *APOE4-WT-P301S* oligodendrocyte DEGs.

Supplementary Table 5 (S5)

DEGs in *APOE4-R47H-P301S* vs. *APOE4-WT-P301S* hippocampal mouse astrocytes. FDR-adjusted *P* values were calculated using MAST. Gene set enrichment analysis hallmark pathways for *APOE4-R47H-P301S* vs. *APOE4-WT-P301S* astrocyte DEGs.

Supplementary Table 6 (S6)

DEGs in *APOE4-R47H/CV* vs. *APOE4-CV/CV* primary microglia at baseline (unstimulated) and after 0N4R tau fibril stimulation. Benjamini–Hochberg-adjusted *P* values reported were calculated using DESeq2. Gene set enrichment analysis hallmark pathways for *APOE4-R47H/CV* vs. *APOE4-CV/CV* primary microglia DEGs at baseline and after tau stimulation. Ingenuity pathway analysis upstream regulator predictions for *APOE4-R47H/CV* vs. *APOE4-CV/CV* tau-stimulated primary microglia DEGs.

